# Could malaria mosquitoes be controlled by periodic releases of transgenic mosquitocidal *Metarhizium pingshaense* fungus? A mathematical modeling approach

**DOI:** 10.1101/2025.05.28.656721

**Authors:** Binod Pant, Etienne Bilgo, Arnaja Mitra, Salman Safdar, Abdoulaye Diabaté, Raymond St. Leger, Abba B. Gumel

## Abstract

Insect pathogenic fungi offer a promising alternative to chemical insecticides for controlling insecticide resistant mosquitoes. One proposed method involves releasing male *Anopheles* mosquitoes contaminated with transgenic *Metarhizium pingshaense* (Met-Hybrid), to lethally infecting females during mating. This study presents a novel deterministic mathematical model to evaluate the impact of this control approach in malaria-endemic areas. The model incorporates two fungus transmission pathways: mating-based transmission and indirect transmission through contact with fungus-colonized mosquito cadavers. We found the fungus cannot establish in the mosquito population without transmission from infected cadavers (in this scenario, the reproduction number of the model is zero). However, if transmission from colonized cadavers is possible, the fungus can persist in the local mosquito population when the reproduction number exceeds one. Simulations of periodic releases of infected male mosquitoes, parameterized using Met-Hybrid-exposed mosquito data from Burkina Faso, show that an 86% reduction in the local female mosquito population can be achieved by releasing 10 Met-Hybrid-exposed male mosquitoes per wild mosquito every three days over six months. This matches the efficiency of some genetic mosquito control approaches. However, a 90% reduction in the wild mosquito population requires, for instance, daily releases of the fungal-treated mosquitoes in a 6:1 ratio for about 5 months, proving less efficient than some genetic approaches. This study concludes that fungal programs with periodic releases of infected males may complement other methods or serve as an alternative to genetic-based mosquito control methods, where regulatory, ethical, or public acceptance concerns restrict genetically-modified mosquito releases.

## 1 Introduction

Vector-borne diseases transmitted by mosquitoes, ticks, sandflies, etc. account for about 17% of all human infectious diseases annually, leading to over 249 million infections and 700,000 deaths worldwide each year [1]. Mosquitoes, labeled as the “world’s deadliest animal” by the United States Centers for Disease Control and Prevention (CDC) [2], are responsible for several human diseases, including malaria, dengue, yellow fever, encephalitis, and filariasis [3]. Malaria stands out as the most important vector-borne disease due to its severe human morbidity and mortality [1]. According to the World Health Organization (W.H.O.) there were an estimated 263 million malaria cases and 597,000 deaths across 85 malaria-endemic countries in 2023 [4]. Additionally, the W.H.O. reported that nearly half of the global population is at risk of dengue, with an estimated 100–400 million infections and 20,000 to 40,000 deaths occurring annually [5]. Consequently, controlling vector-borne diseases, particularly those spread by mosquitoes, is of critical global health importance.

The control of mosquito-borne diseases largely hinges on curbing mosquito populations. Various interventions have been employed to control mosquito populations and reduce the transmission of mosquito-borne diseases. These strategies typically focus on destroying or reducing mosquito breeding habitats and targeting immature and adult mosquitoes with insecticides. Notably, the extensive use of insecticide-based interventions, such as indoor residual spraying (IRS) and long-lasting insecticidal nets (LLINs) [6, 7] has significantly reduced malaria burden in endemic regions. These successes have spurred numerous global initiatives, such as the Global Technical Strategy of the W.H.O., which aims to reduce malaria incidence and mortality in endemic areas by 90% by 2030, and the Zero by 40 initiative, which seeks to eradicate malaria by 2040 [6, 8–10]. Unfortunately, the widespread use of chemical insecticides has led to widespread *Anopheles* resistance to all the chemical compounds used in IRS and LLINs. This resistance, along with reported *Plasmodium* resistance to *artemisinin*-based therapy [11–15], poses significant challenges to the global malaria control and eradication objectives. Numerous alternative biological methods for controlling malaria mosquitoes, such as sterile insect technology (SIT), releasing suitable gene drives, *Wolbachia*-based intervention, vaccines (including RTS, S and R21) and introducing natural predators like fish that target mosquito larvae, have been proposed and some implemented to significantly reduce the population abundance of malaria mosquitoes in endemic areas [16–23].

*Entomopathogenic* fungi are also being considered as an alternative mechanism for control of *Anopheles* mosquitoes. Scholte *et al*. [24] demonstrated that naturally occurring fungi, such as *Metarhizium anisopliae*, which are produced commercially and used against several agricultural insect pests worldwide, show promise as a biological control agent against adult malaria mosquitoes. Specifically, in a field study in Tanzania, hanging *M. anisopliae*-impregnated cotton sheets in houses reduced the lifespan of wild *Anopheles gambiae* mosquitoes that came into contact with them. However, natural *Metarhizium* strains are slow to kill mosquitoes, typically taking 1-2 weeks to reach > 90% mortality [25]. Transgenic *Metarhizium pingshaense* strains have been engineered to express insecticidal toxins like Hybrid (a calcium/potassium channel blocker) and AalT (a sodium channel blocker), which significantly increases their effectiveness against malaria vector mosquitoes [25, 26]. Specifically, the transgenic *M. pingshaense* strain expressing the Hybrid toxin (abbreviated as Met-Hybrid) achieved over 80% mortality within one week, with an average 80% lethal time (LT80) of 5.18, 5.54 and 5.25 days for *An. coluzzii, An. gambiae* and *An Kisumu*, respectively [27]. Transgenic *M. pingshaense* fungi kill mosquitoes through a multi-step process: fungal spores attach to and penetrate the mosquito’s cuticle/exoskeleton, then proliferate inside the body while producing insecticidal toxins like Hybrid and AaIT, ultimately leading to host death due to a combination of tissue damage, nutrient depletion and toxin accumulation. Eventually, the fungus emerges from the dead mosquito and produces new spores on the cadaver surface, ready to infect other mosquitoes [25, 26]. It is important to note that once fungi are engineered with transgenic traits, these genetic modifications are integrated into the fungal genome and can be passed on to their offspring during asexual reproduction, allowing for the potential local production of transgenic fungi in malaria-endemic regions [25, 28].

Similar to SIT and *Wolbachia*-based interventions, which require the periodic release of large numbers of sterile and *Wolbachia*-infected male mosquitoes, respectively, into the natural environment during peak female mosquito abundance, the periodic releases of *M. pingshaense* fungus-exposed male mosquitoes has demonstrated potential in suppressing local wild mosquito populations. This is because the fungus-exposed males can transmit the fungus to the female mosquitoes during mating. However, unlike SIT and *Wolbachia*-infection based strategies, which have proven successful in reducing mosquito-borne diseases in real-world field trials, the transgenic fungal approach is still in its early stages, with results confined to laboratory and semi-field studies [25, 26]. Moreover, while numerous modeling studies have evaluated the impacts of SIT [21,29–35] and *Wolbachia* infection [22,36–38] strategies, no such modeling study has yet been conducted to theoretically assess the potential population-level impact of the transgenic *M. pingshaense* fungus strategy. Thus, to the authors’ knowledge, this study represents the first attempt to theoretically analyze the impact of the fungus-infection-based mosquito control strategy, using a mechanistic mathematical modeling framework.

It should be noted that, in addition to the periodic releases of fungus-exposed male mosquitoes, transgenic fungi can also be deployed in households (although this mechanism is not explored in the current study) and fungus can be transmitted to the mosquitoes through contact with the cadaver of a fungus-infected mosquito (although transmission of fungus through contact with infectious cadaver has been observed in a closed space lab setting, it has not been observed in semi-field setting) [25]. Consequently, the model to be developed in this study focuses on the transmission of the transgenic *M. pingshaense* to uninfected female mosquitoes through contact with fungus-infected males during mating and during contact with cadavers carrying sporulating the fungus.

The paper is structured as follows. Section 2 formulates a baseline model to assess the impact of transgenic *M. pingshaense* fungus, which involves releasing male mosquitoes carrying *M. pingshaense*, on the population abundance of mosquitoes in Burkina Faso. Section 2.1 provides a detailed description and derivation of the baseline values of the parameters, specific to the fungal strain Met-Hybrid and *Anopheles* species *An. coluzzii* in Burkina Faso, used in model simulations. In Section 2.2, the local asymptotic stability property of the model is theoretically assessed. Section 3 extends this baseline *M. pingshaense* model to incorporate the periodic releases of fungus-exposed male mosquitoes. Finally, Section 4 summarizes the main results and provides concluding remarks.

## 2 Formulation of Baseline Model: Single Fungal Application

The baseline model to be formulated in this study is for assessing the population-level impact of a single fungal application (specifically, an initial release of adult male *Anopheles* mosquitoes contaminated with *Metarhizium pingshaense*) on controlling the malaria mosquitoes in Burkina Faso. To formulate the model, the total mosquito population at time *t*, denoted by *N* (*t*), is subdivided into three groups: the total population of male mosquitoes (denoted by *N*_*M*_ (*t*)), unmated female mosquitoes (denoted by *N*_*uF*_ (*t*)) and mated female mosquitoes (denoted by *N*_*rF*_ (*t*)), such that:

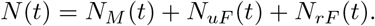

Each of these sub-populations (*N*_*X*_, where the subscript *X* = *M, uF*, or *rF* respectively, represents the population of male, unmated female and mated female mosquitoes in the community) is further divided into mutually-exclusive compartments of mosquitoes susceptible to exposure to *M. pingshaense* fungus (*S*_*X*_ (*t*)), mosquitoes exposed to the fungus with with the fungus on their cuticle/exoskeleton (*E*_*X*_ (*t*)), mosquitoes infected by the fungus with a reduced amount of fungus on their cuticle compared to those in the *E*_*X*_ compartment (*I*_*X*1_(*t*)), and mosquitoes that cannot fly or mate due to the infection, but do not have fungus on their exoskeleton (*I*_*X*2_(*t*)), so that

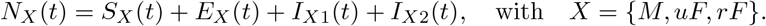

Mosquitoes in the exposed class (*E*_*X*_) are capable of transmitting fungus during mating. However, over time, newly-exposed mosquitoes lose the fungus present on their cuticle. This happens for two reasons: (a) fungal spores penetrate the mosquito cuticle and enter the body, eventually causing illness that leads to the mosquito’s progression to the *I*_*X*1_ and then to the *I*_*X*2_ class [39]; and (b) mosquitoes naturally shed spores over time (for example, through repeated contact with various surfaces and by grooming their fungus carrying legs). Consequently, mosquitoes in the *I*_*X*1_ class have a reduced amount of fungus on their exoskeleton compared to those in the *E*_*X*_ class. However, they still transmit fungus during mating. These progress to *I*_*X*2_ class, which are too sick to fly or mate due to fungal infection.

Furthermore, let *D*(*t*) represent the total population of mosquitoes killed by fungus present in the environment at time *t*. Mosquitoes in the *D*(*t*) compartment do not have fungus on their exoskeleton and, thus, are not able to transmit the fungus to susceptible mosquitoes upon contact. A few days after death, fungus erupts through the cuticle and sporulate, thus forming fungus-carrying infectious cadavers (denoted by the *C*(*t*) compartment) [40].

Let *Ñ*_*rF*_ (*t*) represent the total population of adult female mosquitoes that can lay viable eggs (which will hatch and mature into new adult mosquitoes) at time *t*. Hence,

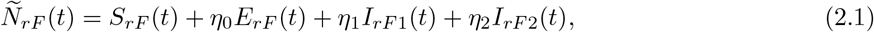

where the modification parameters *η*_0_, *η*_1_ and *η*_2_ (with 0 *≤ η*_2_ *< η*_1_ *< η*_0_ *≤* 1) respectively account for the assumed reduction in the ability of eggs produced by adult fungus-exposed female in the *E*_*rF*_, *I*_*rF*1_, and *I*_*rF* 2_ compartments to develop into adults, compared to eggs laid by mated fungus-free adult female (i.e., mosquitoes in the *S*_*rF*_ compartment). The population of susceptible male mosquitoes (*S*_*M*_) is increased due to the production of new adults at the logistic proliferation rate. It is assumed all new adult mosquitoes are susceptible (male or female), as no vertical transmission of fungus *M. pingshaense* is known to occur [25])

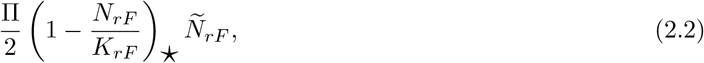

where the factor 1*/*2 accounts for the assumed 50:50 sex ratio, Π is the *per capita* production rate of adult mosquitoes, *K*_*rF*_ is the carrying capacity of the adult mated females in the environment. Although only adult females that can lay viable eggs (*Ñ*_*rF*_) produce new adults, the environmental carrying-capacity for new adults is defined with respect to the total female mosquito population (*N*_*rF*_). The star notation (⋆) below the parenthesis of the logistic proliferation term for new adult mosquitoes, given in (2.2), is used to signify the non-negativity of the logistic eggs oviposition rate defined as 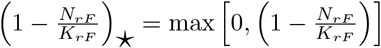.

Furthermore, this population is reduced by exposure to the fungus, due to contact made during mating (at the rate *β*_*sf*_). The term

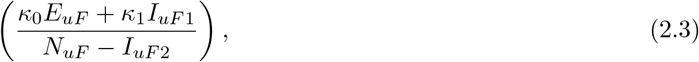

represents the probability that an adult female mosquito capable of mating but not yet mated (i.e., an adult female mosquito that can fly and can mate, specifically, one in the compartments *N*_*uF*_ − *I*_*uF*2_) can transmit the fungus to a susceptible mosquito through contact made during mating (it should be recalled that mosquitoes in the *I*_*uF*2_ compartment are unable to fly or mate). The parameters *κ*_0_ *<* 1 and *κ*_1_ *<* 1, account, respectively, for the assumed reduction in mating ability of fungus-exposed (*E*_*uF*_) and fungus-infected (*I*_*uF*1_) unmated adult females, in comparison to the mating ability of unmated adult females in the *S* compartment (due to their reduced flight capability) [27]. This population is also decreased due to contact with mosquito cadavers that carry sporulating *M. pingshaense* fungus (at the rate *β*_*c*_), and due to the natural death of adult males (mosquitoes in each male compartment are assumed to die naturally at a rate *µ*_*M*_). Thus, the equation for the rate of change of the population of susceptible adult male mosquitoes (*S*_*M*_) is given by:

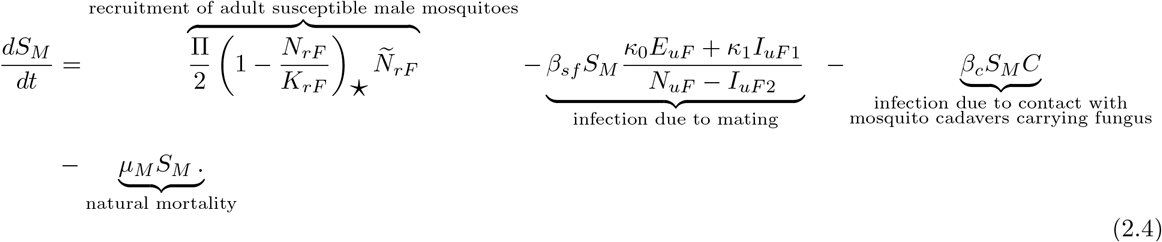

Similarly, the population of unmated adult female mosquitoes susceptible to exposure to the fungus (*S*_*uF*_) increases at the logistic proliferation rate, given by (2.2). This population is decreased due to mating with susceptible adult male mosquitoes (at a rate *γ*_*s*_). The rate *γ*_*s*_ is multiplied by the term 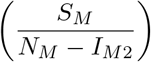, which represents the probability that a mating-capable adult male mosquito (i.e., adult male mosquito in the compartments *N*_*M*_ − *I*_*M*2_) is a fungus-free susceptible mosquito (i.e., the mating is with an adult male mosquito in the *S*_*M*_ compartment). This population further decreases due to exposure to the fungus during mating (at a rate *β*_*sm*_). This rate is modulated by the term 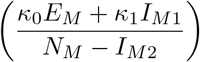, which represents the probability that mating is with a mating-capable adult male mosquito capable of transmitting the fungus during mating. This population is further decreased due to contact with mosquito cadavers carrying sporulating *M. pingshaense* fungus (at the rate *β*_*c*_) and due to the natural death of adult female mosquitoes (mosquitoes in each female compartment are assumed to die naturally at a rate *µ*_*F*_). Thus, the equation for the rate of change of the population of fungus-free (susceptible) adult unmated female mosquitoes (*S*_*uF*_) is given by:

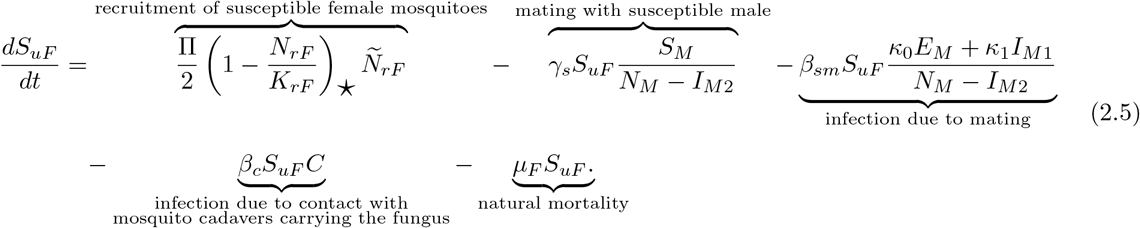

The population of mated adult female mosquitoes susceptible to fungal exposure (*S*_*rF*_) is increased due to the mating of unmated female mosquitoes susceptible to fungal exposure with adult male mosquitoes susceptible to fungal exposure (at the rate 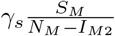). This population is further decreased due to contact with mosquito cadavers that carry *M. pingshaense* (at the rate *β*_*c*_) and by natural death (at the rate *µ*_*F*_). Thus,

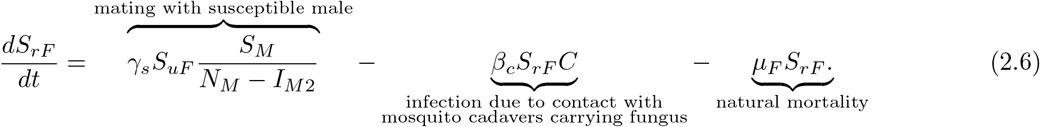

The population of adult male mosquitoes exposed to the fungus (*E*_*M*_) is increased through mating with infectious, mating-capable unmated females (at the rate 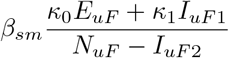), where *N*_*uF*_ − *I*_*uF*2_ is the population of mating-capable adult female mosquitoes. Furthermore, this population is increased due to contact with mosquito cadavers carrying sporulating *M. pingshaense* fungus (at the rate *β*_*c*_). This population is decreased due to progression into the *I*_*M*1_ stage (at the rate *σ*_*E*_) and natural death (at the rate *µ*_*M*_). Thus,

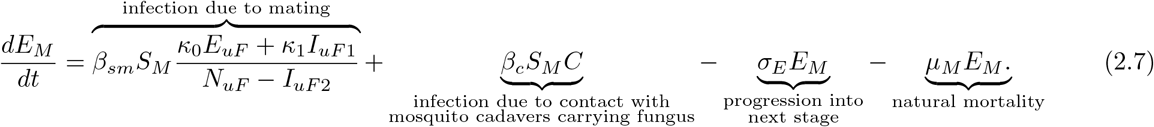

The population of unmated female mosquitoes exposed to the fungus (*E*_*uF*_) is increased due to contact with mosquito cadavers that carry the fungus (at the rate *β*_*c*_). This population is decreased due to mating between unmated fungus-exposed mosquitoes and mating-capable adult male mosquitoes (at the rate 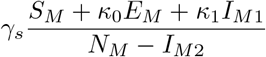, where *N N*_*M*_ − *I*_*M*2_ is the population of mating-capable adult male mosquitoes. Fur-thermore, this population is reduced due to progression into the *I*_*uF*1_ stage (at the rate *σ*_*E*_), and by natural death (at the rate *µ*_*F*_). Thus,

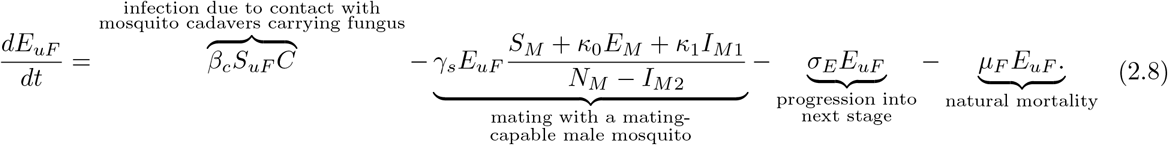

Similarly, the population of mated adult female mosquitoes exposed to the fungus (*E*_*rF*_) is increased due to mating between susceptible unmated adult and male mosquitoes capable of transmitting fungus during mating (at the rate 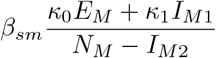). This population is also increased by the mating of exposed unmated female mosquitoes with any mating-capable adult male mosquito (at the rate 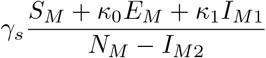). Furthermore, this population is increased due to fungus-exposure caused by contact with mosquito cadavers carrying sporulating fungus (at the rate *β*_*c*_). This population is decreased due to progression to the *I*_*uF*1_ stage (at the rate *σ*_*E*_) and natural death (at the rate *µ*_*F*_). Thus,

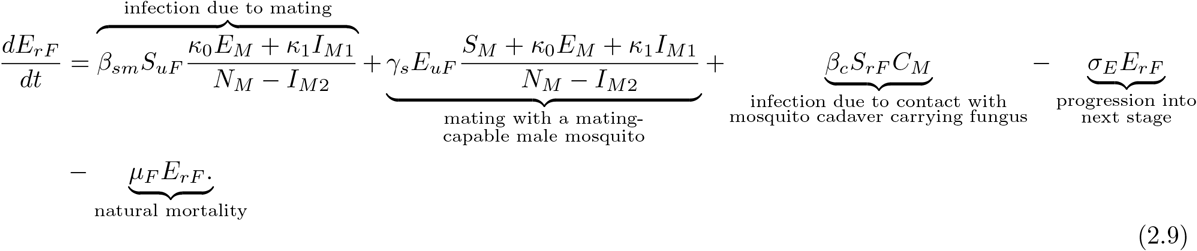

The populations of mosquitoes in the *I*_*M*1_, *I*_*uF*1_, and *I*_*rF*1_ classes are increased due to progression from the *E*_*M*_, *E*_*uF*_, and *E*_*rF*_ classes, respectively (at a rate *σ*_*E*_). These populations are decreased due to progression to the *I*_*M*2_, *I*_*uF*2_, and *I*_*rF* 2_ classes, respectively (at the rate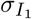) and by fungus-induced mortality (at the rate 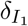). It should be noted that any unmated fungus-infected adult female mosquito in class *I*_*uF*1_ can mate with any mating-capable adult male mosquito (at the rate 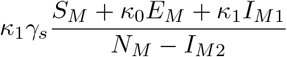), thus transitioning to the class of mated fungus-infected adult female mosquitoes, *I*_*rF*1_. Hence, the equations describing the rate of change of the populations *I*_*M*1_, *I*_*uF*1_, and *I*_*rF*1_ are given, respectively, by:

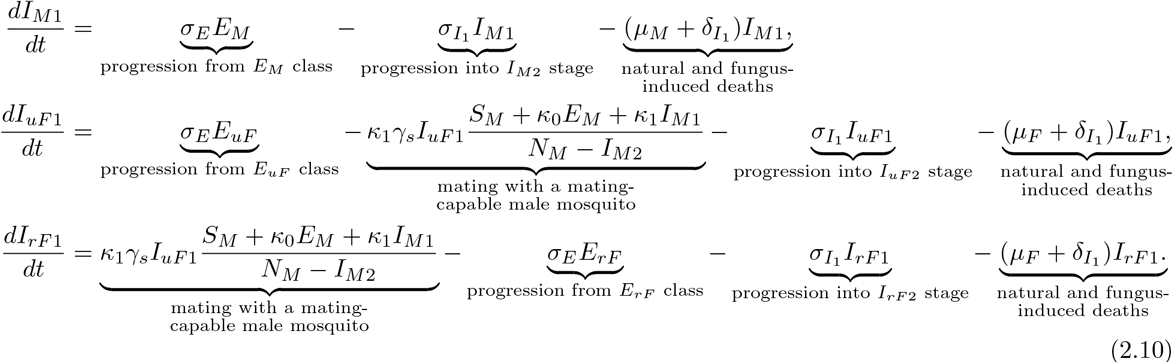

The populations of adult male mosquitoes that cannot fly or mate due to infection with the fungus (*I*_*M*2_), the unmated adult female mosquito that cannot fly or mate due to infection with the fungus (*I*_*uF*2_), and mated adult female mosquito that cannot fly due to infection with the fungus (*I*_*rF* 2_) are increased due to progression from *I*_*M*1_, *I*_*uF*1_ and *I*_*rF*1_ classes, respectively (at the rate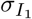). These populations are decreased due to fungus-induced mortality (at a rate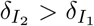). Thus, the equations describing the rate of change of the populations *I*_*M*2_, *I*_*rF* 2_, and *I*_*rF* 2_ are given, respectively, by:

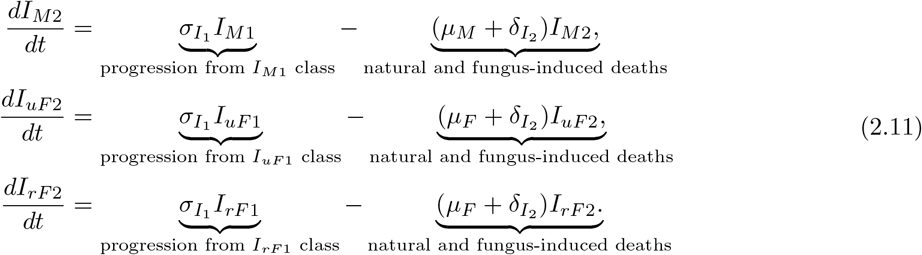

The population of fungus-killed non-infectious mosquitoes (*D*) is increased due to the fungus-induced mortality in the *I*_*M*1_, *I*_*uF*1_ and *I*_*rF*1_ classes (at the rate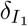) and fungus-induced death in *I*_*M*2_, *I*_*uF*2_, and *I*_*rF* 2_ classes (at the rate 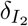). This population is decreased due to the production of infectious fungal spores on cadavers, resulting in infectious cadavers (at the rate *σ*_*D*_) and due to scavenging by organisms such as ants and carrion beetles (at the rate *µ*_*D*_). Thus, the equation for the rate of change of the population *D* is given by:

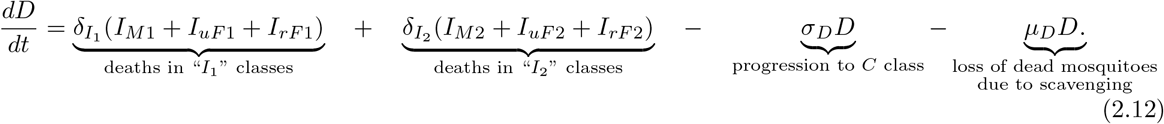

Finally, the number of infectious cadavers (*C*) in the environment increases due to the progression of dead mosquitoes in the *D* compartment to the infectious cadaver compartment *C* (at the rate of *σ*_*D*_). It is decreased due to factors such as decomposition and by scavenging organisms (at the rate *µ*_*C*_). Thus, the equation for the rate of change of the *C* compartment is given by:

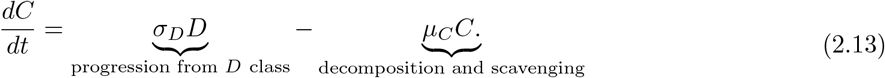

It is convenient to define the following *forces of infection* for fungus transmission due to: (a) mating between a fungus-free susceptible male mosquito with a fungus-infected female mosquito (denoted by *λ*_*sf*_), (b) mating between a fungus-free unmated female mosquito with a fungus-infected male mosquito (denoted by *λ*_*sm*_) and (c) direct contact with infectious mosquito cadavers carrying sporulating fungus (*λ*_*C*_), given respectively, by:

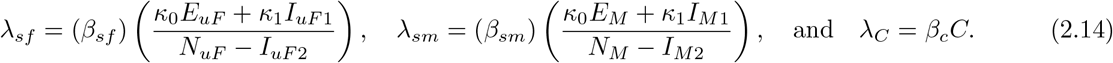

Based on the above derivations and formulations, the baseline model (with a single fungal release) for the transmission dynamics of fungal infection in the adult mosquito population is given by the following deterministic systems of differential equations (where a dot represents differentiation with respect to time *t*):

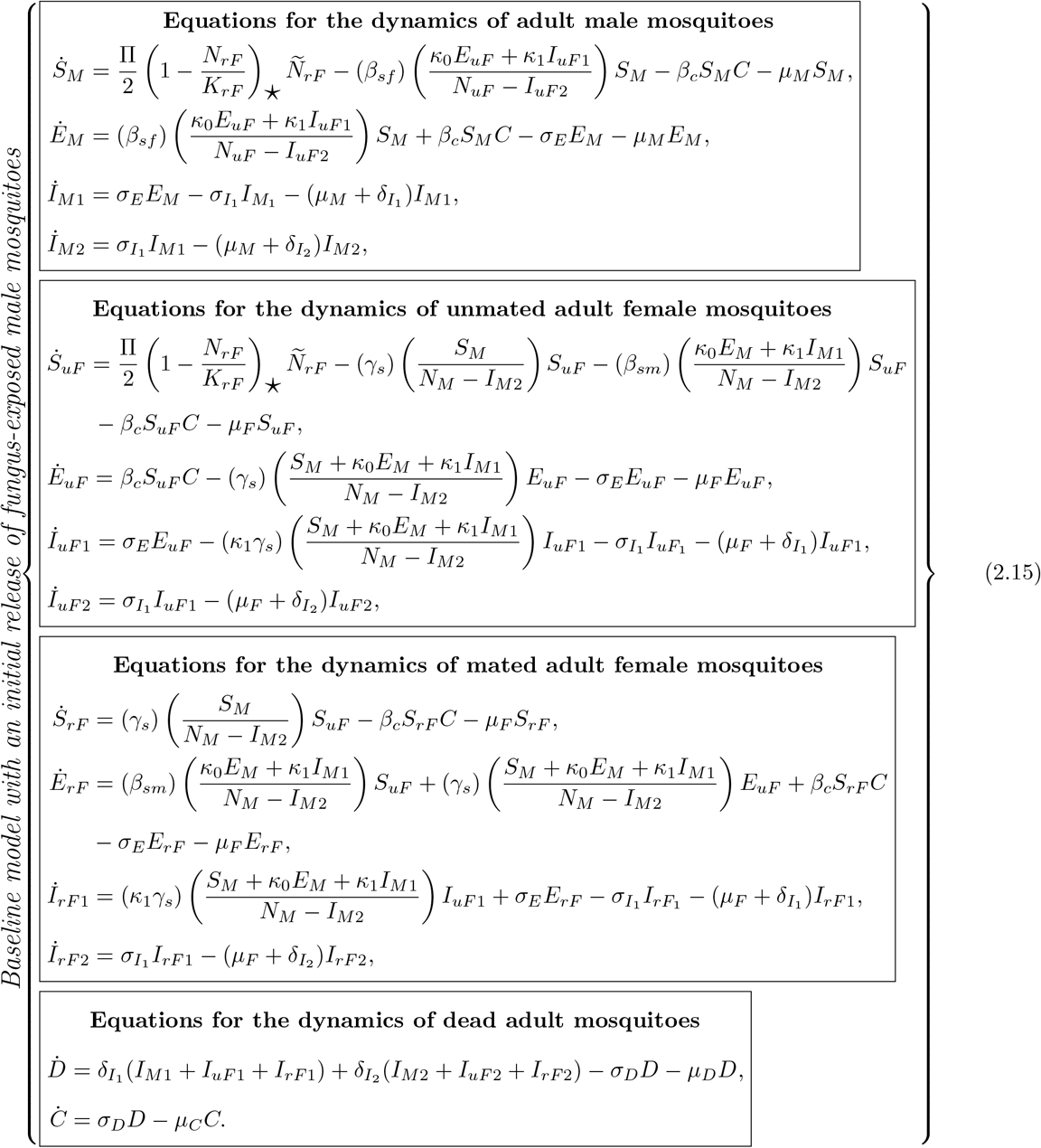

The flow diagram of the baseline model (2.15) is depicted in Figure 2, while the state variables and parameters of the model (2.15) are described in Tables 1 and 2, respectively.

**Table 1:**
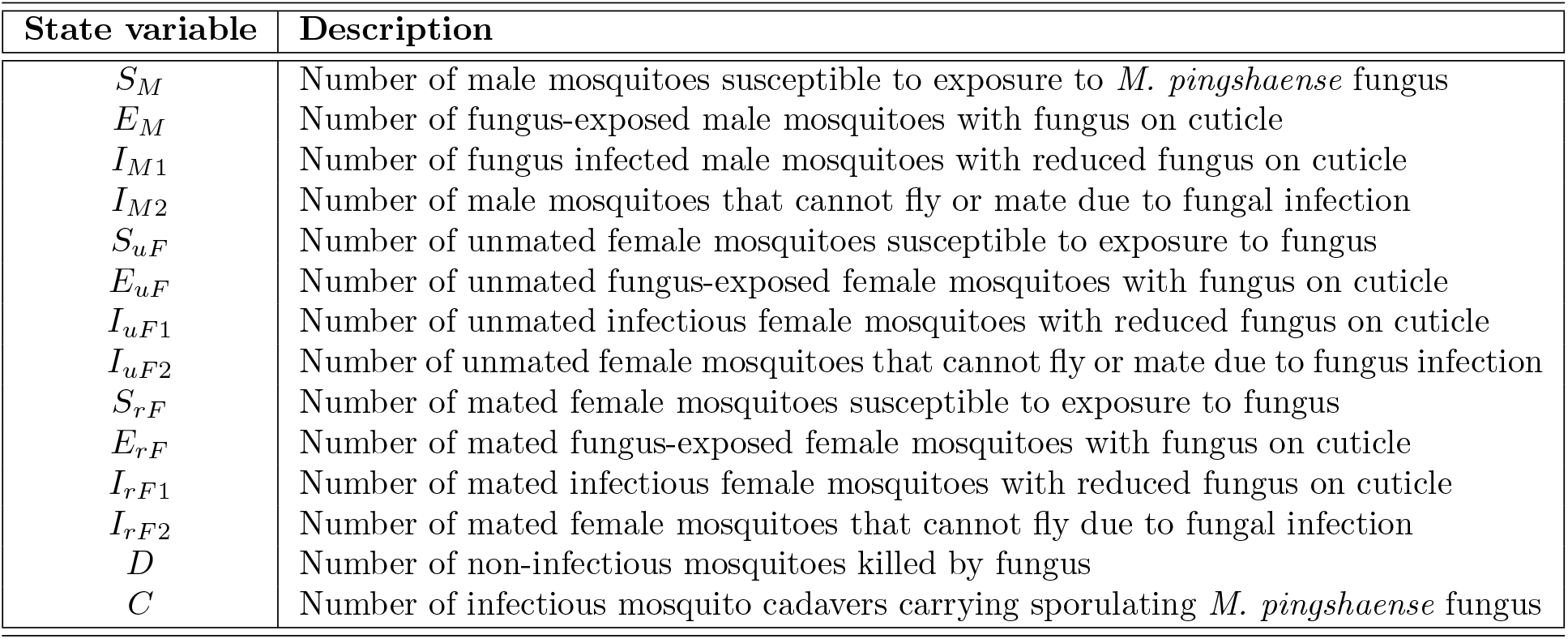
Description of the state variables of the baseline model (2.15).

**Table 2:** Description of the parameters of the baseline model (2.15).

| Parameter | Description |
| --- | --- |
| $\Pi$ | *Per capita* production rate of adult mosquitoes |
| $K_{rF}$ | Carrying capacity of mated female mosquitoes in the environment |
| $\mu_M(\mu_F)$ | Natural death rate of male (female) mosquitoes |
| $\eta_0, \eta_1, \eta_2$ | Modification parameters (with $0 \leq \eta_2 < \eta_1 < \eta_0 \leq 1$) that respectively account for any decrease in the ability of eggs produced by female fungus-infected mosquitoes in the $E_{rF}$, $I_{rF1}$, and $I_{rF2}$ compartments to develop into adults |
| $\sigma_E$ | Progression rate from $E_M$, $E_{uF}$ and $E_{rF}$ to $I_{M1}$, $I_{uF1}$ and $I_{rF1}$, respectively |
| $\sigma_{I_1}$ | Progression rate from $I_{M1}$, $I_{uF1}$ and $I_{rF1}$ to $I_{M2}$, $I_{uF2}$ and $I_{rF2}$, respectively |
| $\gamma_s$ | Mating rate of susceptible unmated female mosquito with susceptible male mosquitoes |
| $\beta_{sf}$ | The rate at which susceptible male mosquitoes acquire fungal infection from mating with an unmated female mosquito capable of transmitting fungus on contact |
| $\beta_{sm}$ | The rate at which susceptible unmated female mosquitoes acquire fungus infection from mating a male mosquito capable of transmitting fungus on contact |
| $\kappa_0$ | Modification parameter (with $0 \leq \kappa_0 \leq 1$) that accounts for the assumed reduction in the mating ability of fungus-exposed mosquitoes in $E_M$ and $E_{uF}$ compartments |
| $\kappa_1$ | Modification parameter (with $0 \leq \kappa_1 \leq 1$) that accounts for the assumed mating reduction in the ability of fungus-infected mosquitoes in $I_M$ and $I_{uF1}$ compartments |
| $\delta_{I_1}(\delta_{I_2})$ | Fungus-induced mortality rate of mosquitoes in the $I_{M1}$, $I_{uF1}$ and $I_{rF1}$ ($I_{M2}$, $I_{uF2}$ and $I_{rF2}$) compartments |
| $\sigma_D$ | Rate at which non-infectious fungus-killed mosquitoes sporulate to become infectious cadaver carrying fungus |
| $\beta_c$ | Rate of infection upon contact with a cadaver carrying sporulated fungus |
| $\mu_D$ | Rate at which non-infectious fungus-killed mosquitoes ($D$) are lost due to scavenging |
| $\mu_C$ | Rate at which infectious cadavers ($C$) are lost due to decomposition and scavenging |

**Figure 1:**
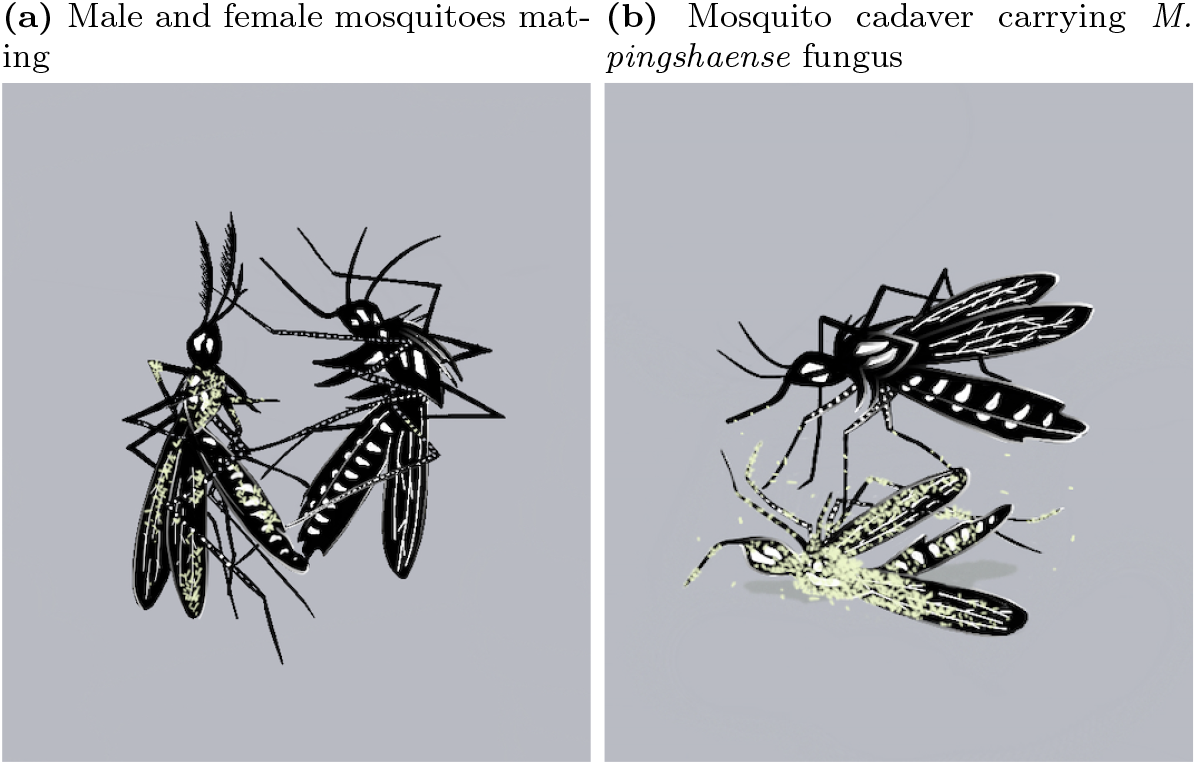
Modes of *M. pingshaense* transmission in mosquitoes. **(a)** *M. pingshaense* transmission to a mosquito of the opposite sex during mating. This sub-figure illustrates a male mosquito with fungal spores on its exoskeleton (left) mating with a susceptible (i.e., fungus-free) female mosquito (right). **(b)** Transmission of *M. pingshaense* through contact with cadavers of dead mosquitoes that carry the fungus. This sub-figure depicts a fungus-susceptible mosquito encountering a cadaver carrying the sporulating *M. pingshaense* fungus.

**Figure 2:**
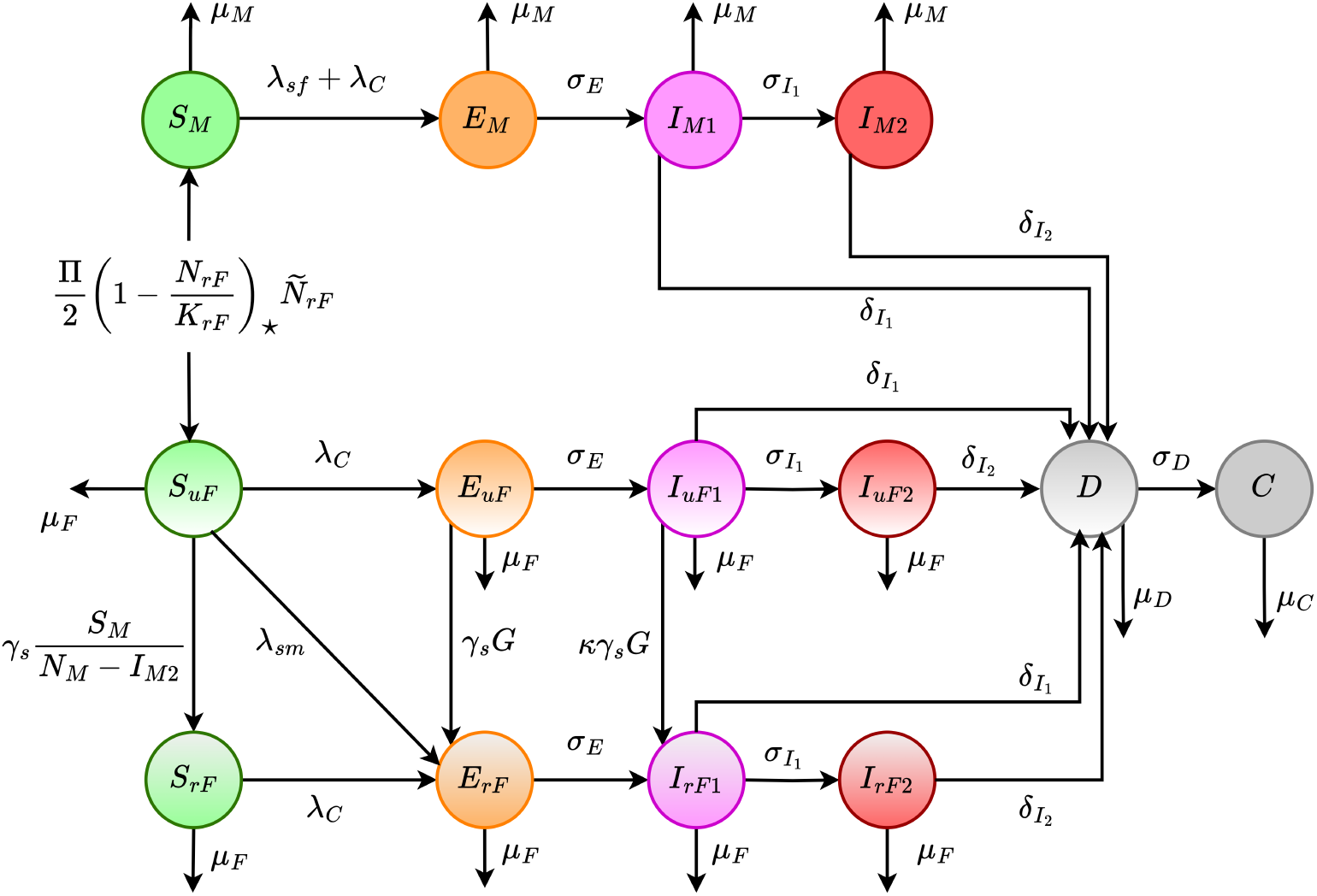
Flow diagram of the baseline model (2.15). The infection rates *λ*_*sf*_, *λ*_*sm*_ and *λ*_*C*_ are defined in Equation (2.14) while *Ñ*_*rF*_ is defined in Equation (2.1). For simplicity in the flow diagram, we define 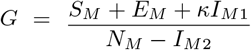.

Some of the main assumptions made in the formulation of the baseline model with a single fungal release (2.15) are (*inter alia*):

i. It is assumed that an initial baseline number of fungus-exposed male mosquitoes are present in the environment following a single release [39]. In other words, an initial number of fungus-infected male mosquitoes are introduced into the environment, and this initial release is not repeated. Mathematically, this assumption implies that the initial number of fungus-infected male mosquitoes (*E*_*M*_ (0)) is positive.
ii. The populations of new adult susceptible male and adult unmated susceptible female mosquitoes grow at a logistic proliferation rate, as described by equation (2.2) [9, 15].
iii. Adult female mosquitoes do not mate more than once. This is a reasonable assumption since female mosquitoes of the *Anopheles gambiae* species complex, the primary vectors of malaria, typically mate only once during their lifetime under field conditions [41, 42]. Moreover, *Anopheles coluzzii*, the vector that will be used to parametrize the model (2.15), belongs to this complex, and adult females of this species also mate only once [43].
iv. *M. pingshaense* is transmitted to the mosquito via two main mechanisms: (a) mating (b) contact with infectious mosquito cadavers carrying sporulating fungus. Although fungal spores (the infectious propagule) are only produced after mosquito death, many cadavers are scavenged before sporulation, limiting the secondary transfer of fungus through cadavers [25].
v. For simplicity, it is assumed that *M. pingshaense* is not capable of living saprophytically in the soil. While soil-dwelling strains exist [44, 45], it is reasonable to assume that a mosquito cadaver carries too little fungus for the fungus to establish in soil.

It should be noted that the model formulated in this section is for the transmission dynamics of *M. pingshaense* fungus in the mosquito population. In the subsequent sections, the model will be parameterized and simulated with respect to a transgenic *M. pingshaense* (Met-Hybrid), which kills mosquitoes faster and at lower spore doses than wild-type strains [27].

### 2.1 Baseline values of the parameters of the baseline model

In this section, the numerical values of the parameters of the baseline model (2.15) will be discussed or justified, in the context of the Met-Hybrid fungal strain and the *An. coluzzii* species prevalent in Burkina Faso. The average lifespan of adult male (1*/µ*_*M*_) and female (1*/µ*_*F*_) *An. coluzzii* mosquitoes is approximately 2 weeks [46]. Hence, we set *µ*_*M*_ = 1*/*14 *per* day and *µ*_*F*_ = 1*/*14 *per* day, respectively. The *per capita* production rate of new adult mosquito recruitment (Π) can be defined as:

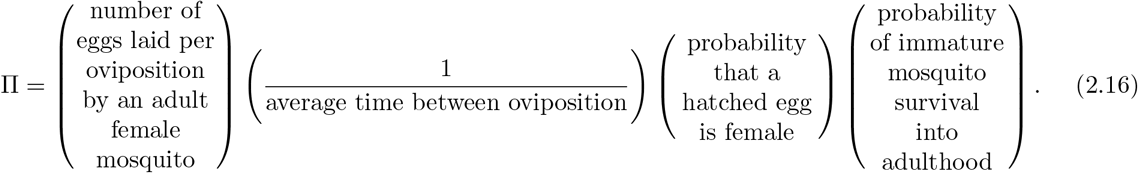

Adult female mosquitoes are estimated to lay between 100 to 300 eggs per oviposition [47, 48] every third night [49]. Assuming a 50:50 male-to-female sex ratio, the probability of a hatched egg being female is 1/2. According to the American Mosquito Control Association Public Health Pest Control [50], the probability of an immature mosquito surviving the aquatic stages (egg, larval, and pupal) to become an adult is less than 5%. Consequently, it is reasonable to set the number of eggs laid per oviposition to be 200 and the probability of immature mosquito survival to adulthood at 3%. Hence, 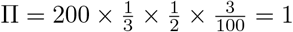 *per* day.

Data reported by Lovett et al. [26] indicates that adult female mosquitoes do not experience a reduction in their ability to produce offspring during the fungus-exposed period, that is, the first 2.5 days of exposure to the fungus. Consequently, the modification parameter that accounts for the reduction in egg production ability in adult female mosquitoes within the *E*_*rF*_ class (*η*_0_) is set to 1. The study by Lovett et al. [26] further revealed that while 77.8% of uninfected (control) adult female mosquitoes laid eggs, only 24.7% of those infected with Met-Hybrid did so (see Figure S6 in the Supplement of [26]). This suggests that the modification parameter for the reduction in egg laying ability of fungus-infected adult female mosquitoes (*η*_1_) is 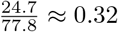. Thus, *η*_1_ = 0.32. Additionally, for simplicity, it is assumed that mated female mosquitoes unable to fly (i.e., those in the *I*_*rF* 2_ compartment) do not lay eggs that hatch and mature into adult mosquitoes. This assumption is reasonable, as mosquitoes that cannot fly may encounter difficulties in reaching breeding sites for egg-laying. Therefore, the modification parameter accounting for the reduction in the quality of eggs laid by adult female mosquitoes in the *I*_*rF* 2_ compartment (*η*_2_) is set to zero. That is, *η*_2_ = 0.

The parameter *σ*_*E*_, which represents the progression from *E*_*M*_, *E*_*uF*_ and *E*_*rF*_ classes to the *I*_*M*1_, *I*_*uF*1_ and *I*_*rF*1_ classes, is derived from the empirical studies by Bilgo et al [27, 51], which showed that Met-Hybrid takes 2 to 3 days (with an average of 2.5 days) to penetrate the host cuticle. Thus, *σ*_*E*_ = 1*/*2.5 *per* day. To estimate the parameter *σ*_*I*_, which pertains to the progression rate of mosquitoes from the flying-capable fungus-infected state (i.e., mosquitoes in the *I*_*M*1_, *I*_*uF*1_, and *I*_*rF*1_ compartments) to the flying-incapable fungus-infected state (i.e., *I*_*M*2_, *I*_*uF*2_, and *I*_*rF* 2_), mosquito flight data will be utilized. Specifically, Bilgo *et al*. [27] observed a significant decline in the flight ability of fungus-exposed mosquitoes starting on day 3 post exposure. We assume that, on average, a fungus-infected mosquito in the *I*_*M*1_, *I*_*uF*1_, and *I*_*rF*1_ compartments remains flight-capable for 1 day before losing the ability to fly. Consequently, the progression rate *σ*_*I*_ is set at 1 *per* day.

A fungus-susceptible (i.e., fungus-uninfected) mosquito that mates with a mosquito carrying fungus on its exoskeleton will acquire fungus spores during mating and eventually succumb to the infection [39]. In other words, the probability of transmission of fungus during mating is nearly 100%. Adult female mosquitoes typically mate within 2-3 days (with a mean of 2.5 days) from emergence to adulthood [52], leading to a mating rate of unmated susceptible female mosquitoes with susceptible male mosquitoes (*γ*_*s*_) of 1/2.5 *per* day. Thus *γ*_*s*_ = 1*/*2.5 *per* day. Similarly, adult male mosquitoes mate soon after emergence to adulthood. Therefore, we assume that the rate at which susceptible male mosquitoes acquire fungal infection through mating with a unmated adult female mosquito capable of transmitting the fungus (*β*_*sf*_) is 1/2.5 *per* day.

Hence, *β*_*sf*_ = 1*/*2.5 *per* day. Likewise, the rate at which susceptible unmated adult female mosquitoes acquire a fungal infection through mating with a mating-capable mated male mosquito capable of transmitting the fungus upon contact (*β*_*sm*_) is 1/2.5 *per* day, resulting in *β*_*sm*_ = 1*/*2.5 *per* day. Bilgo *et al*. [27] demonstrated that Met-Hybrid lacks irritant and excito-repellent properties. Thus, it is assumed that mosquitoes exposed to or infected with, the fungus do not deter potential mating partners due to the presence of fungus on their exoskeleton. However, fungal infection can impose a fitness cost by diminishing the mating ability of fungus-infected adult mosquitoes.

The impact of fungal infection on mating ability remains unknown. We deduce the modification parameter value, which accounts for reduced mating ability, from mosquito flight data, as mosquitoes, including *Anopheles coluzzii*, predominantly mate while flying. Mating is initiated by male mosquitoes forming a swarm at dusk, followed by adult female mosquitoes flying into the swarm [53, 54]. Additionally, males are known to detect potential mates in the swarm by recognizing faint female flight tones [55]. According to Table 3 in the empirical study by Bilgo *et al*. [27], both fungus-exposed and non-exposed (control) mosquitoes display similar flight abilities on days 1 and 2, with comparable percentages in flight time. This indicates that, during the exposure period (i.e., while the mosquitoes are in the *E*_*M*_, *E*_*uF*_ and *E*_*rF*_ compartments), their flight capability remains unaffected. Consequently, the reduction in mating ability for fungus-exposed mating-capable mosquitoes in the *E*_*M*_ and *E*_*uF*_ compartments due to fungal exposure (denoted by the parameter *κ*_0_) is 1, meaning there is no reduction. However, Bilgo *et al*. [27] reported a 38% and 74% reduction in flight time on days 3 and 4 post-exposure, respectively, with an average reduction of 56%. Therefore, it is assumed that the reduction in mating for adult mosquitoes in the *I*_*M*1_ and *I*_*uF*1_ compartments (denoted by the parameter *κ*_1_) is 0.56.

**Table 3:** Baseline values of the parameters for the single release model (2.15).

| Parameter | Value | Source |
| --- | --- | --- |
| $\Pi$ | 1 mosquito/day | Computed (using Equation (2.16)) |
| $K_{rF}$ | $10^5$ mosquitoes | Assumed |
| $\mu_M, \mu_F$ | $1/14 \text{ day}^{-1}$ | [46] |
| $\eta_0$ | 1 (dimensionless) | [26] |
| $\eta_1$ | 0.32 (dimensionless) | [26] |
| $\eta_2$ | 0 (dimensionless) | [26] |
| $\sigma_E$ | $1/2.5 \text{ day}^{-1}$ | [27] |
| $\sigma_{I_1}$ | $1 \text{ day}^{-1}$ | [27] |
| $\gamma_s$ | $1/2.5 \text{ day}^{-1}$ | [56] |
| $\beta_{sf}$ | $1/2.5 \text{ day}^{-1}$ | [57] |
| $\beta_{sm}$ | $1/2.5 \text{ day}^{-1}$ | Assumed |
| $\kappa_0$ | 1 (dimensionless) | [27] |
| $\kappa_1$ | 0.56 (dimensionless) | [27] |
| $\sigma_D$ | $1/3.5 \text{ day}^{-1}$ | Assumed |
| $\delta_{I_1}$ | $0.15 \text{ day}^{-1}$ | [26, 27] |
| $\delta_{I_2}$ | $0.07 \text{ day}^{-1}$ | [26, 27] |
| $\beta_c$ | $0 \text{ day}^{-1}$ | [25] |
| $\mu_D$ | $1/7 \text{ day}^{-1}$ | Assumed |
| $\mu_C$ | $1/7 \text{ day}^{-1}$ | Assumed |

**Table 4:**
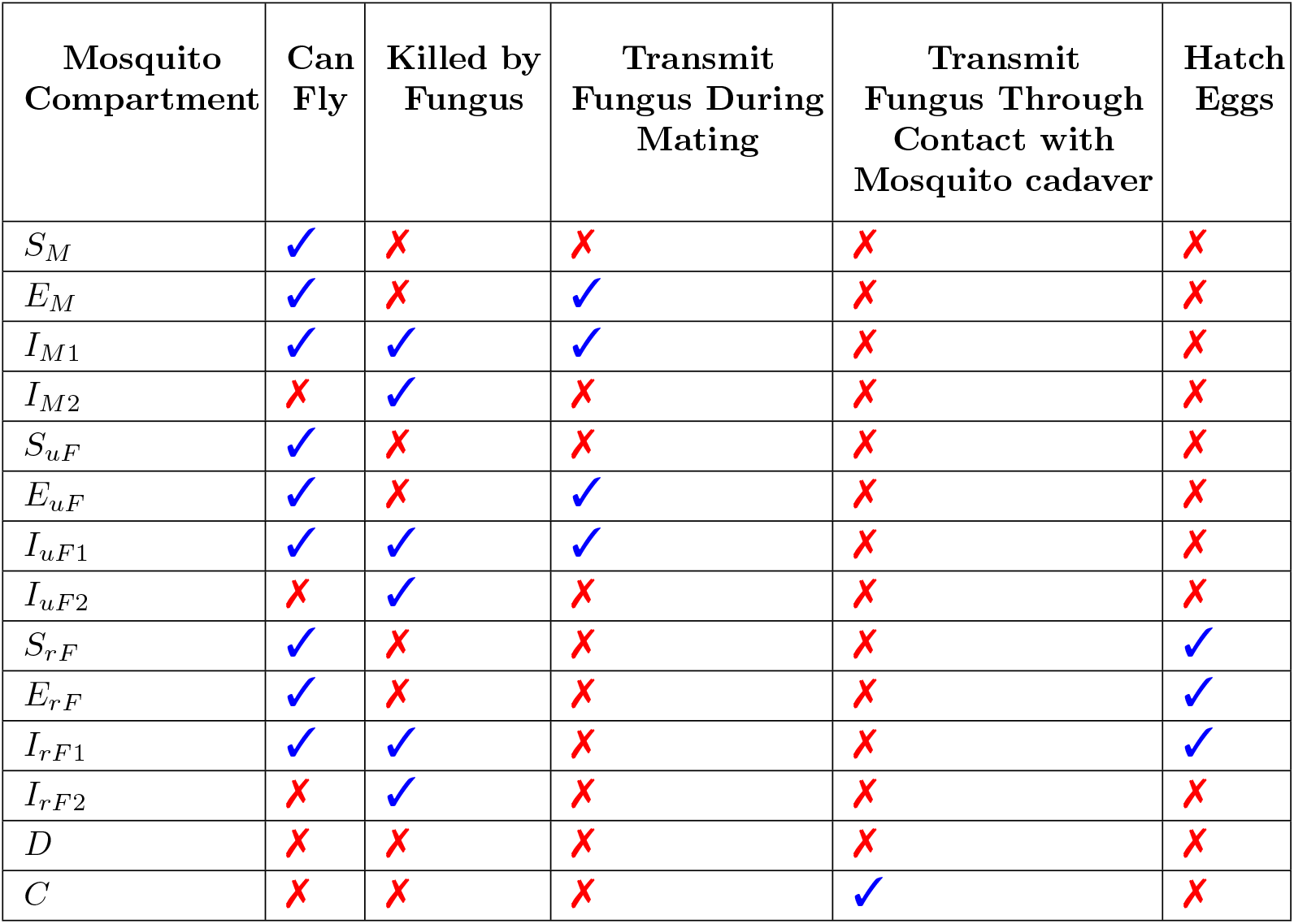
Impact of *M. pingshaense* infection on the ecological status and fitness of mosquitoes, assessed through their flying ability, survival, fungal transmission during mating or contact with cadaver, and oviposition and egg hatching. The fungus’s effect on mosquitoes in different ecological compartments and fungal-exposure stages is evaluated by examining their flight capability, fungus-induced mortality, transmission of fungus during mating, transmission via infectious cadavers, and egg hatching success.

| Mosquito Compartment | Can Fly | Killed by Fungus | Transmit Fungus During Mating | Transmit Fungus Through Contact with Mosquito cadaver | Hatch Eggs |
| --- | --- | --- | --- | --- | --- |
| $S_M$ | ✓ | ✗ | ✗ | ✗ | ✗ |
| $E_M$ | ✓ | ✗ | ✓ | ✗ | ✗ |
| $I_{M1}$ | ✓ | ✓ | ✓ | ✗ | ✗ |
| $I_{M2}$ | ✗ | ✓ | ✗ | ✗ | ✗ |
| $S_{uF}$ | ✓ | ✗ | ✗ | ✗ | ✗ |
| $E_{uF}$ | ✓ | ✗ | ✓ | ✗ | ✗ |
| $I_{uF1}$ | ✓ | ✓ | ✓ | ✗ | ✗ |
| $I_{uF2}$ | ✗ | ✓ | ✗ | ✗ | ✗ |
| $S_{rF}$ | ✓ | ✗ | ✗ | ✗ | ✓ |
| $E_{rF}$ | ✓ | ✗ | ✗ | ✗ | ✓ |
| $I_{rF1}$ | ✓ | ✓ | ✗ | ✗ | ✓ |
| $I_{rF2}$ | ✗ | ✓ | ✗ | ✗ | ✗ |
| $D$ | ✗ | ✗ | ✗ | ✗ | ✗ |
| $C$ | ✗ | ✗ | ✗ | ✓ | ✗ |

Recall that the exposed stage *E*_*X*_ (with *X* = *M, uF, rF*) lasts an average of 2.5 days while the *I*_*X*1_ stage lasts on average 1 day. The percentage of fungus-infected mosquitoes that die on days 2.5, 3 and 3.5 days post inoculation to fungus is 10%, 11%, and 15%, respectively, with an average of 12% [26]. Thus, the probability that a mosquito in *I*_*X*1_ compartment dies due to fungus is (noting that *µ* = *µ*_*M*_ = *µ*_*F*_):

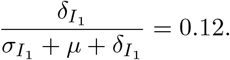

Substituting 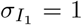 *per* day and *µ* = 1*/*14 *per* day from Table 3, and solving for 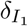, gives 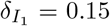 *per* day. Meanwhile, the average percentage of mosquitoes that die 3.5 to 7 days after inoculation with fungus is 40% [26]. Hence, the probability that a mosquito in the *I*_*X*2_ compartment (with *X* = {*M, uF, rF*}) dies due to fungus infection is (noting that *µ* = *µ*_*M*_ = *µ*_*F*_):

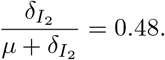

Using *µ* = 1*/*14 *per* day from Table 3, and solving for 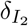, gives 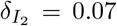 *per* day. Finally, the values of the parameters related to fungus-carrying cadavers are determined as follows. Laboratory observations, indicate that the time for a fungus killed mosquito to begin sporulating (i.e., progression time from D to C compartment) ranges from 4 to 5 days, with a mean of 3.5 days. Therefore, *σ*_*D*_ = 3.5 *per* day. In a semi-field setting, Lovett et al. [25] observed that many cadavers are scavenged before sporulation occurs (i.e., fungus-killed mosquitoes in the *D* compartments are scavenged before they transition to sporulating, fungus-carrying cadavers compartment *C*). Moreover, the chances of mosquitoes, as flying insects with aquatic larval stages, encountering fungus-carrying cadavers is likely minimal. Consequently, the rate of fungus exposure due to contact with mosquitoes in the C compartment (*β*_*C*_) is set to zero. Finally, we assume that the average time for decomposition (1*/µ*_*C*_) and/or scavenging (1*/µ*_*D*_) of dead mosquitoes is approximately seven days (hence, *µ*_*D*_ = *µ*_*C*_ = 1*/*7 *per* day).

### 2.2 Existence and local asymptotic stability of fungus-free equilibria

The baseline model (2.15) has two *fungus-free equilibria (FFE)*, namely a trivial *mosquito-free* FFE (denoted by ℰ_*t*_) and a nontrivial *mosquito-present* FFE (denoted by ℰ_*nt*_), given, respectively, by:

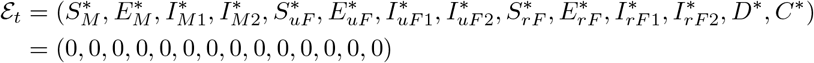

and,

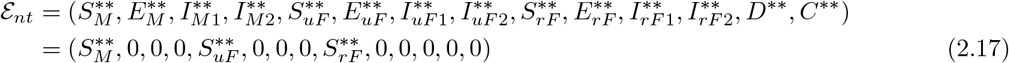

where,

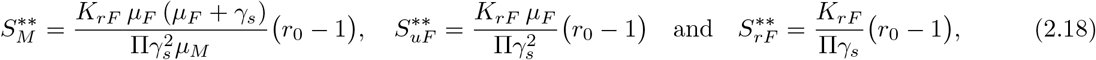

with,

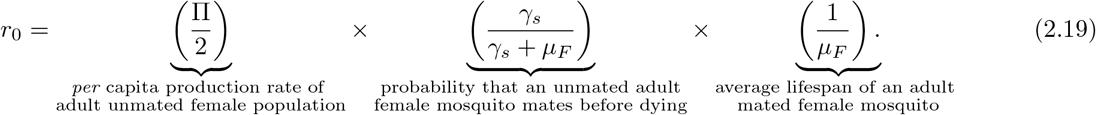

It follows from (2.18) that the trivial equilibrium ℰ_*nt*_ exists if and only if *r*_0_ > 1. Furthermore, when *r*_0_ = 1, the non-trivial mosquito-present FFE (ℰ_*nt*_) collapses into the trivial mosquito-free FFE (ℰ_*t*_). This result is summarized below.

#### Theorem 2.1.

*The baseline model* (2.15) *has a trivial mosquito-free and fungus-free equilibrium (*ℰ_*t*_*) that always exists. The model has a non-trivial mosquito-present and fungus-free equilibrium (*ℰ_*nt*_*) which exists whenever r*_0_ > 1.

The quantity *r*_0_ determines the survival or extinction of the mosquito population since the mosquito population only exists when *r*_0_ > 1, while no mosquito exists for *r*_0_ *≤* 1. The threshold quantity, *r*_0_, is the *net reproduction value for mated adult female mosquitoes*, which has an ecological interpretation of the expected number of mated female mosquitoes produced by a single mated female mosquito over its lifetime [58]. It is a product of the *per capita* production rate of adult female population 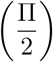, the probability that an unmated adult female mosquito mates before dying 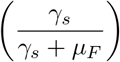 and the average lifespan of an adult female mosquito 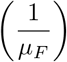. Hence it is evident that a high *per capita* production rate and mating rate increases *r*_0_ while a high mortality rate decreases *r*_0_. In other words, both *per capita* production rate and mating rates need to be high enough to compensate for the mortality rate for the non-trivial mosquito-present equilibrium to exist.

Since the trivial equilibrium (ℰ _*t*_) of the baseline model (2.15) is not ecologically realistic (since mosquitoes are expected to always exist in the natural environment in the malaria-endemic areas), its asymptotic stability properties will not be rigorously analyzed. These properties will only be analyzed for the non-trivial mosquito-present equilibrium (ℰ _*nt*_), as below. The local asymptotic stability of the non-trivial equilibrium (ℰ _*t*_) will be explored using the *next generation operator method* [59,60]. Specifically, using the notation in [60], it follows that the associated non-negative matrix of new fungus infection terms (𝔽) and the M-matrix of all linear transition terms (𝕍) are given, respectively, by

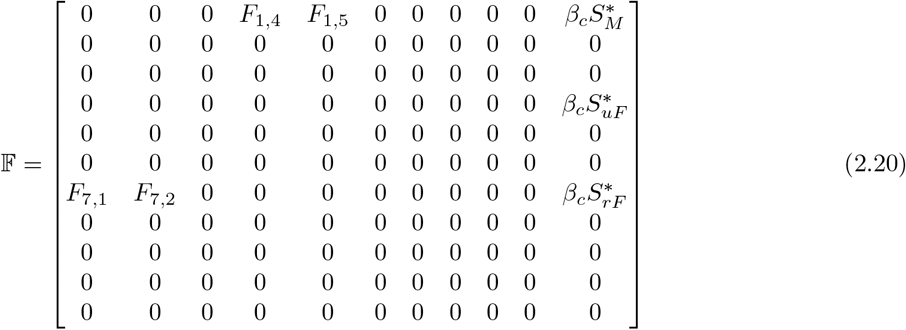

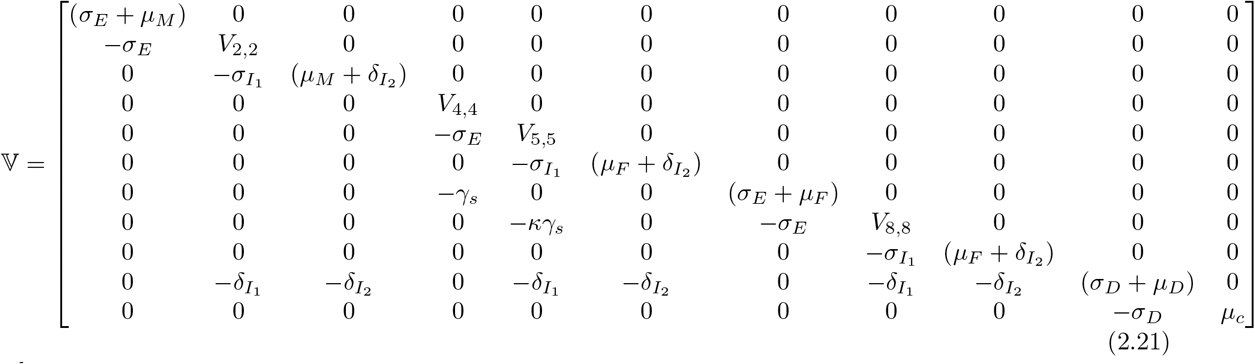

where,

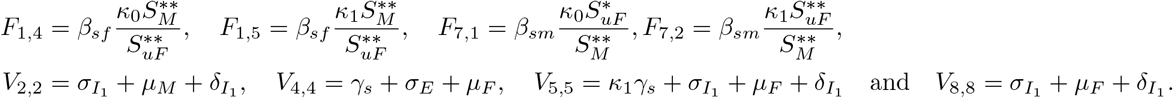

It is convenient to define the following quantity (where *ρ* is the spectral radius):

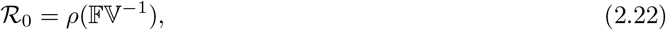

where the matrices 𝔽 and 𝕍 are as given by (2.20) and (2.21), respectively. Note that this reproduction threshold is expressed in this concise manner, since its closed-form expression is too complex to be written here.

The result below follows from Theorem 2 of [60].

#### Theorem 2.2.

*The non-trivial fungus-free equilibrium of the baseline model* (2.15) *(which exists only if r*_0_ > 1*) is locally-asymptotically stable if* ℛ _0_ *<* 1, *and unstable whenever* ℛ _0_ > 1.

The ecological interpretation of Theorem 2.2 is that a small influx of fungus-exposed mosquitoes will not generate a large number of fungus-exposed mosquitoes in the environment when ℛ _0_ *<* 1. In other words, releasing a small number of fungus-exposed mosquitoes when ℛ _0_ *<* 1 will still lead to the elimination of fungus from the mosquito population. Conversely, if ℛ _0_ > 1, releasing even a small number of fungus-exposed mosquitoes may allow fungus to establish within the mosquito population.

The quantity ℛ _0_ is the *basic reproduction number*, which is defined as the average number of fungus-exposed male mosquitoes generated by a male mosquito with fungus on its exoskeleton. A similar definition of ℛ _0_ also exists with respect to female mosquitoes with fungus on their exoskeleton.

#### 2.2.1 Special case of the baseline model with no transmission of fungus from mosquito cadavers

Although the baseline model (2.15) considers the transmission of fungus to mosquitoes through contact with fungus-infected mosquito cadavers, there is no definitive evidence supporting this transmission mechanism [25]. In this section, we will evaluate the impact of relaxing the assumption for fungal transmission *via* contact with fungus-infected cadavers. Specifically, we will theoretically assess a special case of the baseline model (2.15) where the parameter for fungus transmission from infectious cadavers, *β*_*c*_, is set to zero.

Let 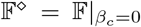. In this special case of the baseline model (2.15) with *β*_*c*_ = 0, it can be shown that the spectral radius of the matrix 𝔽^⋄^𝕍^−1^ is zero, as all eigenvalues of the aforementioned 𝔽^⋄^𝕍^−1^ matrix are zero. In other words, when fungal transmission from infectious cadavers does not occur (i.e., when *β*_*c*_ is set to zero in the baseline model), the associated reproduction number (defined as 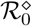) is precisely zero. This is because, without cadaver-mediated transmission, fungus-exposed male mosquitoes cannot generate new fungus-exposed males. In this scenario, fungal spread is *unidirectional* (e.g., male-to-female *via* mating only; Figure 3(a)). For ℛ _0_ > 0, a *cyclical* transmission loop is required, where the fungus must complete a transmission pathway that generates new fungus-exposed males. Infectious cadavers serve as critical *intermediate* components in these fungal transmission pathways, enabling the fungus to cycle back to susceptible male mosquitoes through either male-to-cadaver-to-male transmission or male-to-female-to-cadaver-to-male transmission. These two fungal transmission pathways are further illustrated through the following specific scenarios:

**Figure 3:**
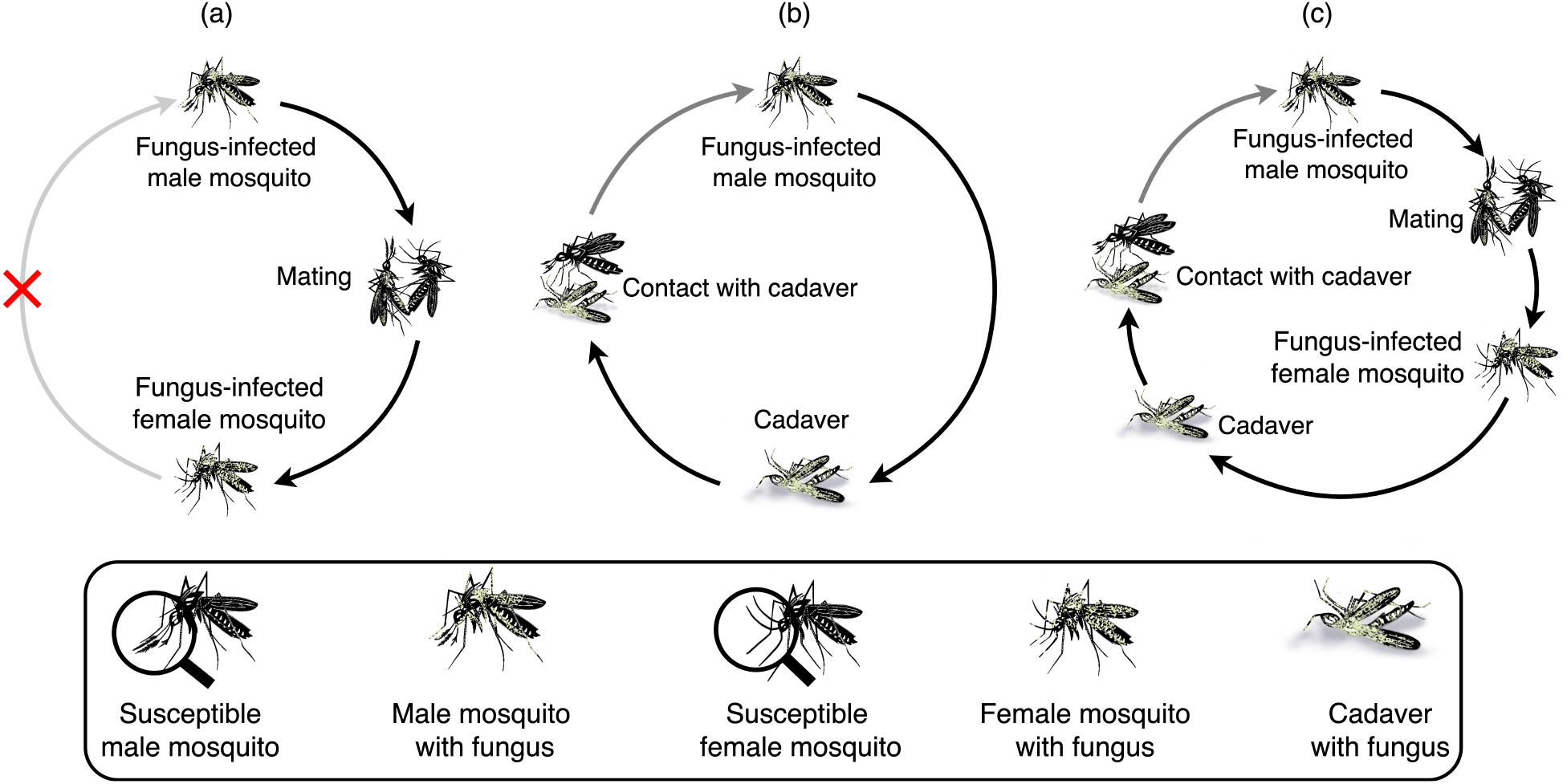
Illustration of why fungal transmission through cadavers (*β*_*c*_ > 0) is critical to complete the cyclical transmission loop, where fungus-exposed or fungus-infected male mosquitoes generate new fungus-exposed or fungus-infected male mosquitoes. (a) A fungus-infected male mosquito capable of mating (i.e., *E*_*M*_ or *I*_*M* 1_) mates with a fungus-susceptible unmated female mosquito (*S*_*uF*_), resulting in a fungus-exposed mated female mosquito (*E*_*rF*_). This mated female mosquito cannot generate additional fungus-infected male mosquitoes because, once mated, female mosquitoes do not mate again. (b) A fungus-infected male mosquito (i.e., *E*_*M*_, *I*_*M* 1_ or *I*_*M* 2_) eventually dies and becomes an infectious cadaver (*C*). When a susceptible male mosquito (*S*_*M*_) comes in contact with this cadaver (*β*_*c*_ > 0), it becomes fungus-exposed (*E*_*M*_), which, with disease progression, can eventually turn into *I*_*M* 1_ and *I*_*M* 2_. (c) A fungus-infected male mosquito capable of mating (i.e., *E*_*M*_ or *I*_*M* 1_) mates with a susceptible unmated female mosquito (*S*_*uF*_), resulting in a fungus-exposed mated female mosquito (*E*_*rF*_), which eventually dies and becomes an infectious cadaver (*C*). When a susceptible male mosquito (*S*_*M*_) comes in contact with this cadaver (*β*_*c*_ > 0), it becomes fungus-exposed (*E*_*M*_), and can eventually turn into *I*_*M* 1_ and *I*_*M* 2_.

i. **Fungus-infected male** *→* **cadaver** *→* **fungus-infected male transmission pathway**: A male mosquito infected with fungus (i.e., mosquitoes in the *E*_*M*_, *I*_*M*1_ or *I*_*M*2_ class) may eventually succumb to the fungal infection, resulting in an infectious cadaver (i.e., mosquito in the *C* class). A susceptible male mosquito that encounters this infectious cadaver can contract the infection (see Figure 3(b)). This newly-infected male mosquito (in the *E*_*M*_ class) can later transition to the *I*_*M*1_ and *I*_*M*2_ classes, which can progress to form infectious cadavers, thereby perpetuating the male mosquito-cadaver-male mosquito fungal transmission pathway.
ii. **Fungus-infected male** *→* **female-infected** *→* **cadaver** *→* **fungus-infected male transmission pathway:** When a fungus-exposed (*E*_*M*_) or mating-capable fungus-infected (*I*_*M*1_) male mosquito mates with a fungus-free susceptible female mosquito (*S*_*uF*_), the susceptible mosquito becomes a mated fungus-exposed female mosquito, which can eventually succumb the fungal infection and turn into an infectious cadaver (*C*) carrying sporulating fungus on its exoskeleton. If a susceptible male mosquito encounters this cadaver, it can contract the fungal infection, becoming a fungus-exposed male mosquito (see Figure 3(c)).

##### Remark.

*A special case of the baseline model* (2.15) *with the non-existent transmission of fungus from cadavers carrying fungus (i*.*e*., *β*_*c*_ = 0*) has an associated reproduction number of zero (i*.*e*., 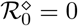).

We claim the following result for the special case of the baseline model with *β*_*c*_ = 0.

##### Lemma 2.1.

*Assume that the solution of the baseline model* (2.15) *is non-negative. The baseline model without fungal transmission by infectious cadavers (i*.*e*., *β*_*c*_ = 0*) has no fungus-present equilibrium*.

*Proof*. Consider the baseline model with no fungal transmission by infectious cadavers (i.e., consider the model (2.15) with *β*_*c*_ = 0). Setting *β*_*c*_ = 0 into the equation for *Ė*_*uF*_ gives:

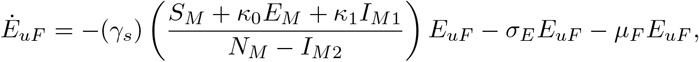

from which it follows that

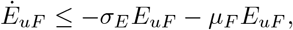

so that (using the integrating factor 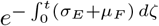

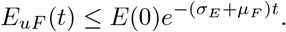

Since the right-hand side of the above inequality decays exponentially to zero for positive parameters, and we assumed *E*_*uF*_ (*t*) is non-negative, we have

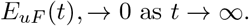

Recall that the equation for the rate of change of the *I*_*UF*1_ population is given by:

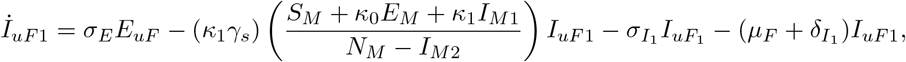

which implies

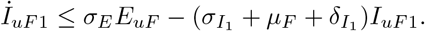

so that (using an integrating factor):

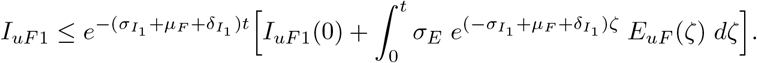

Since 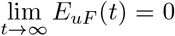, and we assumed *I*_*uF*1_(*t*) to be non-negative, it follows that 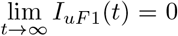. Using the same approach, it can be seen that the rest of the fungi-infected compartments of the model tend to zero as *t* approaches infinity. That is,

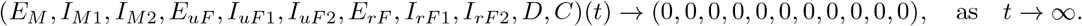

□

The ecological implication of Lemma 2.1 is that *M. pingshaense* fungus cannot independently survive or persist in the mosquito population over the long term without fungal transmission through infectious cadavers. Therefore, for the case of the model with *β*_*c*_ = 0, an external mechanism, such as the periodic mass release of fungus-exposed male mosquitoes, is necessary to sustain fungal presence in the mosquito population. Additionally, the theoretical results in this section also highlight the importance of employing various metrics, beyond just the reproduction number, to realistically assess the population-level impact of the fungal-based mosquito control program.

Intuitively, in ecological situations where adult female mosquitoes mate frequently, it is plausible that a fungus-exposed male mosquito could contribute to the generation of other fungus-exposed male mosquitoes, thereby completing the cycle depicted in Figure 3(a). Mathematically-speaking, if adult female mosquitoes mate frequently, there would be no need to divide the adult female mosquito population into unmated and mated adult female mosquitoes. Instead, it would suffice to examine the transmission dynamics of the fungus among the adult mosquito population, divided into adult male and adult female groups. These types of models are known to have the basic reproduction number expressed as the geometric mean of the constituent reproduction numbers associated with each of the two groups, resulting in an overall nonzero basic reproduction number. If the basic reproduction number exceeds one, the fungus could potentially be sustained within the mosquito population. Studies have shown that adult female mosquitoes of species such as *Aedes albopictus* (also known as Asian tiger mosquito) [61] and *Anopheles freeborni* (also known as western malaria mosquito) [62] can mate multiple times in their lifetime. However, this is not the case with the *Anopheles coluzzii*, the malaria vector prevalent in the study area in Burkina Faso considered in this study [43].

## 3 Model with Periodic Releases of Fungus-exposed Male Mosquitoes

Entomologists in Burkina Faso, including co-authors Bilgo and Diabaté, have recently conducted empirical studies on the impact of releasing fungus-infected male mosquitoes in both laboratory and semi-field settings in this West African nation. Shortly after emerging from pupae to adulthood, the mosquitoes are separated according to sex to obtain a cohort of virgin males, which are then exposed to Met-Hybrid, the transgenic *M. pingshaense* strain. One method for exposing these virgin male mosquitoes to the fungus involves placing them inside a cage lined with fungus-sprayed cloth. As the male mosquitoes encounter the cloth, they become fungus-exposed mosquitoes. This method of exposure may require modification to enhance efficiency when scaling up for larger deployments of fungus-exposed male mosquitoes.

The baseline model (2.15) is now extended to account for the periodic release of fungus-exposed adult male mosquitoes into the environment. This entails defining [21, 63],

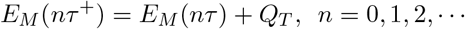

where *τ* is the time lag between successive release of fungus-exposed adult male mosquitoes and *nτ* ^+^ is the moment immediately after the *n*th release. At each time *t* = *nτ*, a constant number of fungus-exposed adult male mosquitoes (*Q*_*T*_) are released into the environment. Similarly, the other state variables (denoted by *Y*) are redefined as:

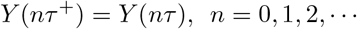

where *Y* = {*S*_*M*_, *I*_*M*1_, *I*_*M*2_, *S*_*uF*_, *I*_*uF*_, *I*_*uF*_, *S*_*rF*_, *I*_*rF*_, *I*_*rF*_, *C, D*}. The resulting periodic male release model is given by the impulsive deterministic system in Equation (A.1) of Appendix A.

### 3.1 Study site for the periodic fungus-exposed male mosquito release

The study site for the periodic male release is Soumousso, a savanna town in Burkina Faso, with a total human population of 118,402 [64] (see Figure 4 for the map of Soumousso, and its location within Burkina Faso). Soumousso hosts five species of malaria vectors that are prevalent throughout the year: *An. coluzzii, An. gambiae, An. funestus, An. nili*, and *An. arabiensis*, with *An. gambiae* being the predominant species year-round [65].

**Figure 4:**
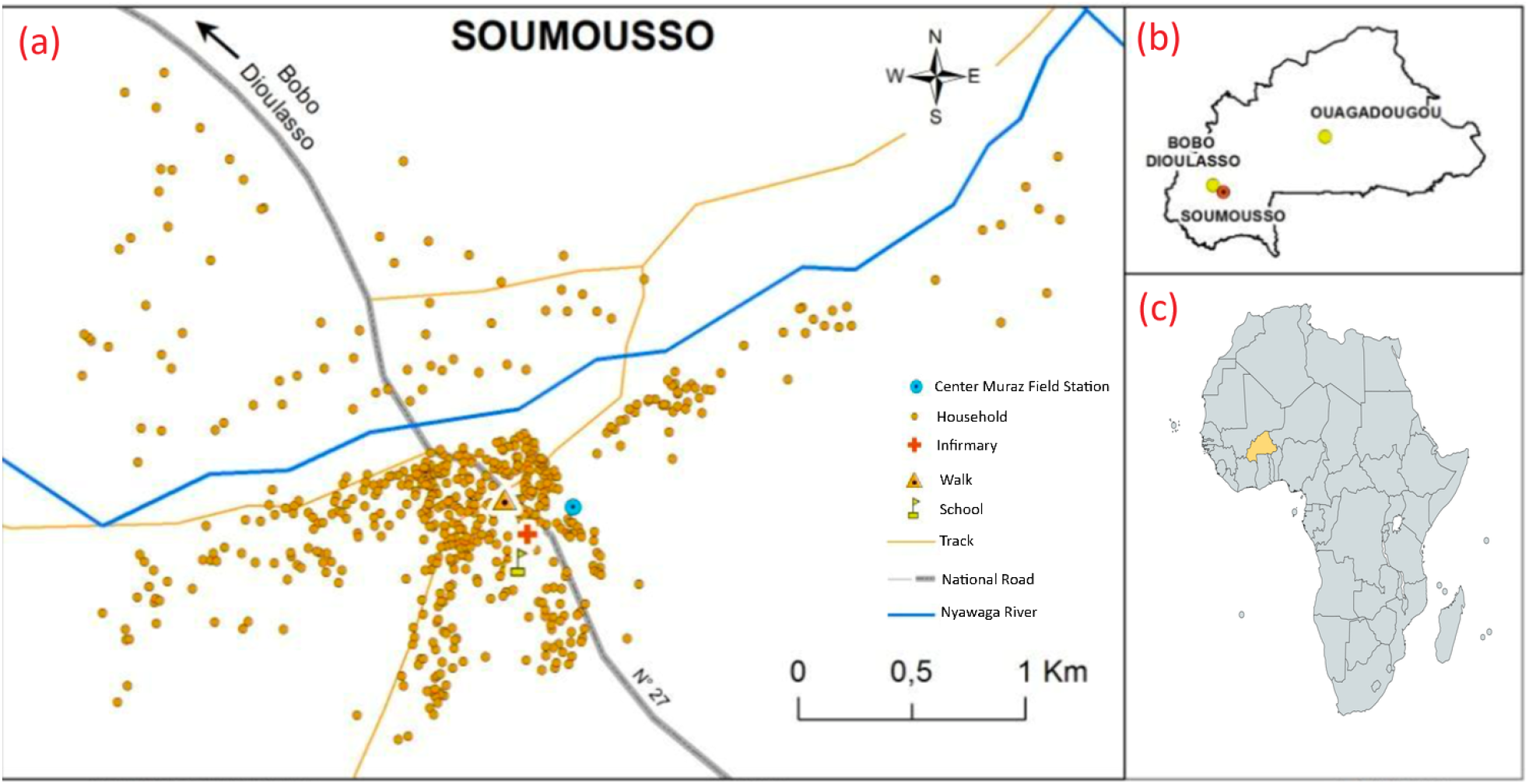
Overview of the study location, its surrounding geography, and its context within the country of Burkina Faso and the African continent. **(a)** Map of the study site, Soumousso, a savanna town in Burkina Faso. **(b)** Map of Burkina Faso depicting the relative location of the study site, Soumousso, in relation to two major cities in Burkina Faso, namely Bobo Dioulasso (40 kilometers away) and Ouagadougou (361 kilometers away). **(c)** Map of Africa with Burkina Faso highlighted in yellow. The map of Soumousso is adapted from [65].

### 3.2 Simulations of the extended model with periodic fungus-exposed male releases

In this section, the baseline model with periodic male releases, given by the equations in (A.1), where *Q*_*T*_ is the average number of fungus-exposed male mosquitoes that are periodically released every *τ* units of time, is simulated to assess the potential impact of the periodic releases of fungus-exposed male mosquitoes on the population abundance of the wild mosquitoes in the study area. For simplicity, we assume that infectious cadavers do not transmit the fungus to susceptible mosquitoes (i.e., we set *β*_*c*_ = 0). Before simulating the periodic release model (A.1), we first simulated the baseline single-release model (2.15) for one year, to allow the system to settle to a mosquito-present steady-state. The periodic male release was then initiated. Let *N*_*F*_ (*t*) = *N*_*uF*_ (*t*) + *N*_*rF*_ (*t*) be the total number of adult female mosquitoes at the study site at time *t*, where *N*_*uF*_ (*t*) and *N*_*rF*_ (*t*) are the total number of unmated and mated female population at time *t*. Let 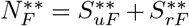 be the total number of female mosquitoes at the mosquito-present fungus-free equilibrium (FFE), where 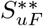 and 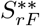, defined in (2.18), are the numbers of unmated and mated female susceptible populations at the non-trivial mosquito-present fungus-free equilibrium, respectively.

The impact of a release program, with specific release frequency *τ* and release amount *Q*_*T*_, is assessed by accounting for the reduction in the total number of adult female mosquito population at time *t* (*N*_*F*_ (*t*)), relative to the total number of adult female mosquitoes at the mosquito-present FFE 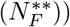. Specifically, the quantity 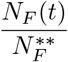 represents the fraction by which the adult female mosquito population has been reduced, relative to the FFE 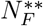. Hence, we define the *percentage reduction* (*P*_*R*_) as the percentage of adult female mosquitoes reduced, in comparison to the baseline value at the FFE 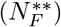. More formally, *P*_*R*_ is given by:

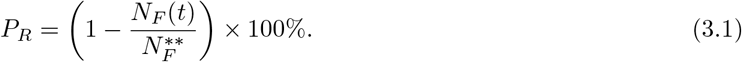

The target percent reduction is set at 90%, aligning with the United Nations’ Sustainable Development Goal to decrease the global malaria burden by 90% [8] (other studies employing SIT and similar mosquito bio-control measures, such as *Wolbachia*-infection based methods [66, 67], have also set 90% as the target reduction for the malaria vector population).

The periodic release model (A.1) is initially simulated using the baseline parameter values in Table 3 for the scenario where 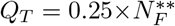 fungus-exposed male mosquitoes are released daily (i.e., *τ* = 1). The results, illustrated in Figure 5, display the temporal dynamics of various mosquito populations (or compartments) before, during, and after the periodic releases of fungus-exposed male mosquitoes. Specifically, this figure indicates that, although the populations of unmated and mated female mosquitoes fluctuate during the time when the fungus-exposed male mosquitoes are released, such changes are relatively minor, and the mosquito populations (both male and female) quickly revert to their pre-release steady-state once the program is halted. In fact, Figure 6 reveals that the total female mosquito population decreases during the release period of fungus-exposed male mosquitoes; however, only about a 6% reduction in the adult female population was achieved under this program.

**Figure 5:**
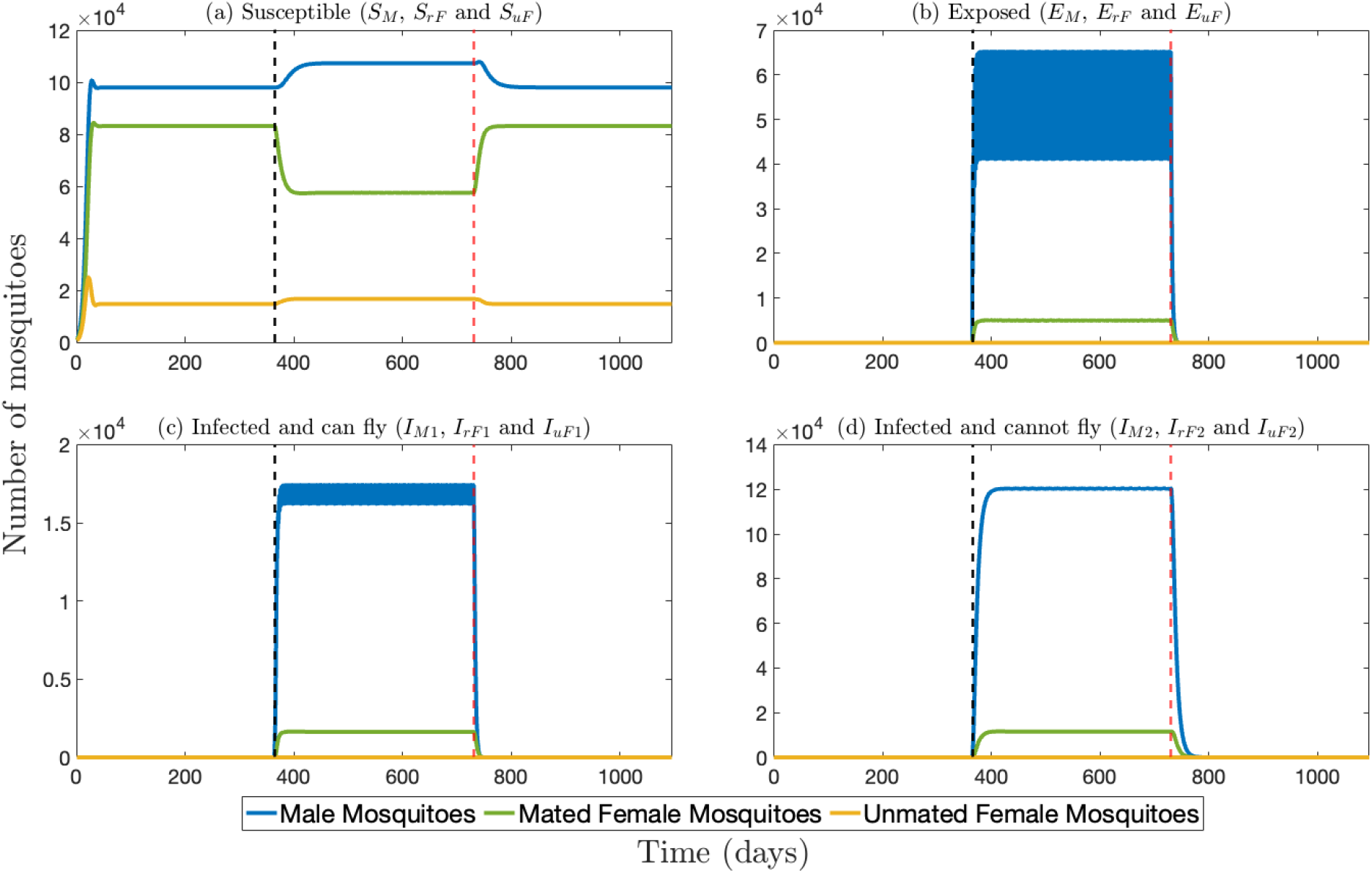
The impact of daily periodic releases of fungi-exposed male mosquitoes on the mosquito population. The periodic release model (A.1) is simulated over three years as follows: initially, the model is run for one year without any releases to allow the system to settle into its mosquito-present steady state; subsequently, fungus-exposed male mosquitoes are released periodically for one year; finally, the model is simulated for an additional year to evaluate the impact of stopping the releases. The four panels depict the compartments for (a) Susceptible, (b) Exposed, (c) Infected and capable of flight, and (d) Infected and unable to fly mosquitoes. In each panel, the populations of male, mated female, and unmated female mosquitoes are denoted by blue, green, and yellow curves, respectively. The vertical dashed black line indicates the start of the periodic releases of fungus-exposed male mosquitoes, and the vertical dashed red line marks when the release stops. The number of mosquitoes released per impulse is 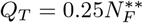. The remaining parameter values are provided in Table 3.

**Figure 6:**
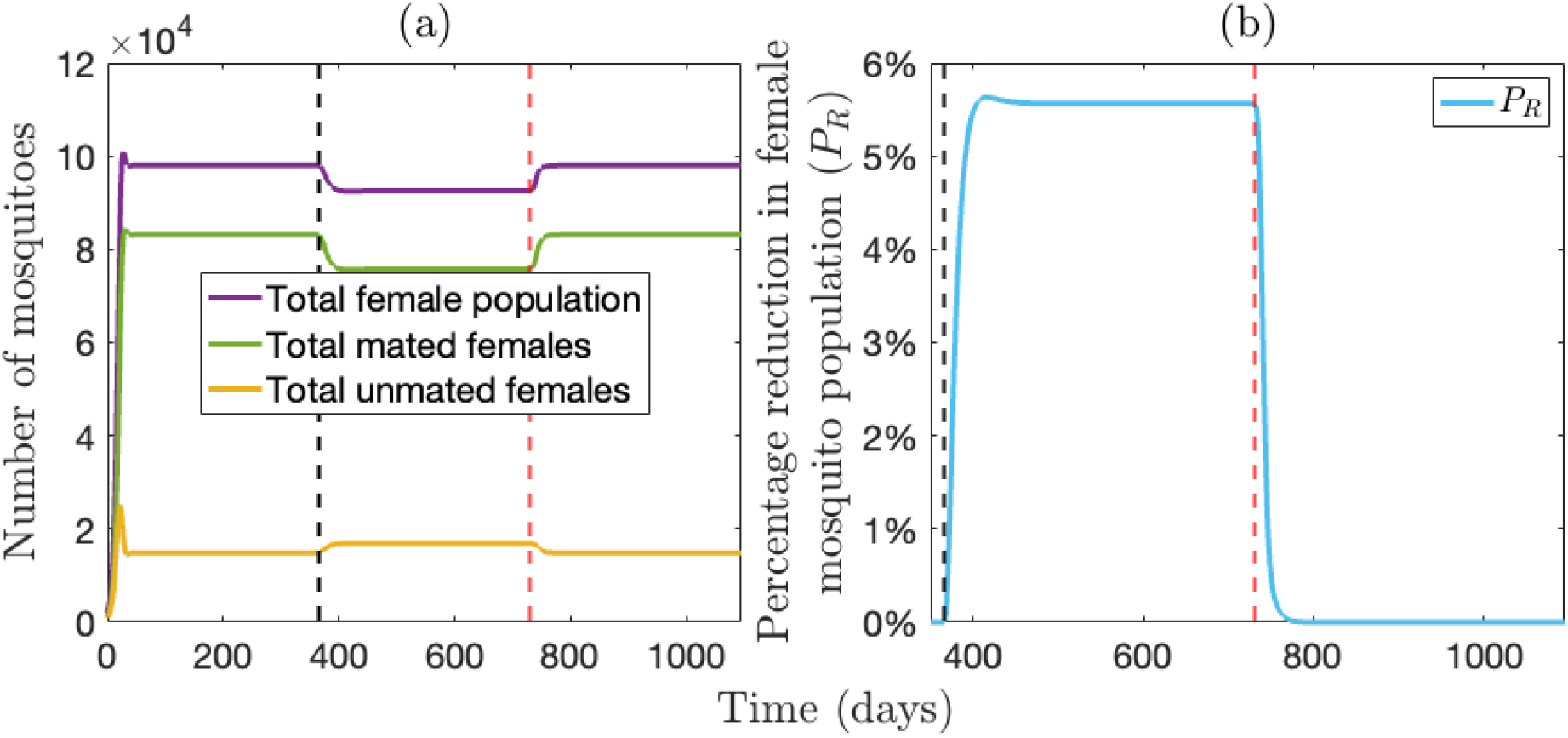
The impact of daily periodic releases of fungi-exposed male mosquitoes on the mosquito population. The periodic release model (A.1) is simulated over three years as follows: initially, the model is run for one year without any releases to allow the system to settle into its mosquito-present steady state; subsequently, fungus-exposed male mosquitoes are released periodically for one year; finally, the model is simulated for an additional year to evaluate the impact of stopping the releases. (a) The change in the total female population, mated female and unmated female population. (b) The percentage reduction in the female mosquito population (*P*_*R*_) due as a function of time. The number of mosquitoes released per impulse is 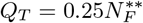. The remaining parameters are provided in Table 3.

Figure 7 presents a contour plot depicting the percent reduction in the total population of adult female mosquitoes (*P*_*R*_, as defined by (3.1)), as a function of the total number of male mosquitoes released *per* release period (*Q*_*T*_) and various release frequencies (*τ*), which correspond to different durations of the male release program. For example, Figures 7(a)-(d) demonstrate that achieving 90% reduction in the total number of adult female mosquitoes, compared to their population size at the pre-release steady-state, is unattainable if the male release program is conducted for only one to about 4 months. This holds true regardless of the quantity of fungus-exposed male mosquitoes released *per* release period (*Q*_*T*_) and release frequency (*τ*) considered in the figure. However, Figures 7(e)-(i) reveal that the 90% reduction can be achieved if the program is ran for at least 5 months with large numbers (e.g., 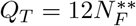) of fungi-infected males being released frequently (e.g., daily). Given the assumption of a 50:50 sex ratio in the wild mosquito population, this 12:1 release ratio (i.e., releasing 12 Met-Hybrid-exposed male mosquitoes for every wild female mosquito at the pre-fungal steady state) is equivalent to releasing exposed males in a 6:1 ratio (i.e., 6 exposed males for every wild mosquito in the local environment at a pre-fungal steady state).

**Figure 7:**
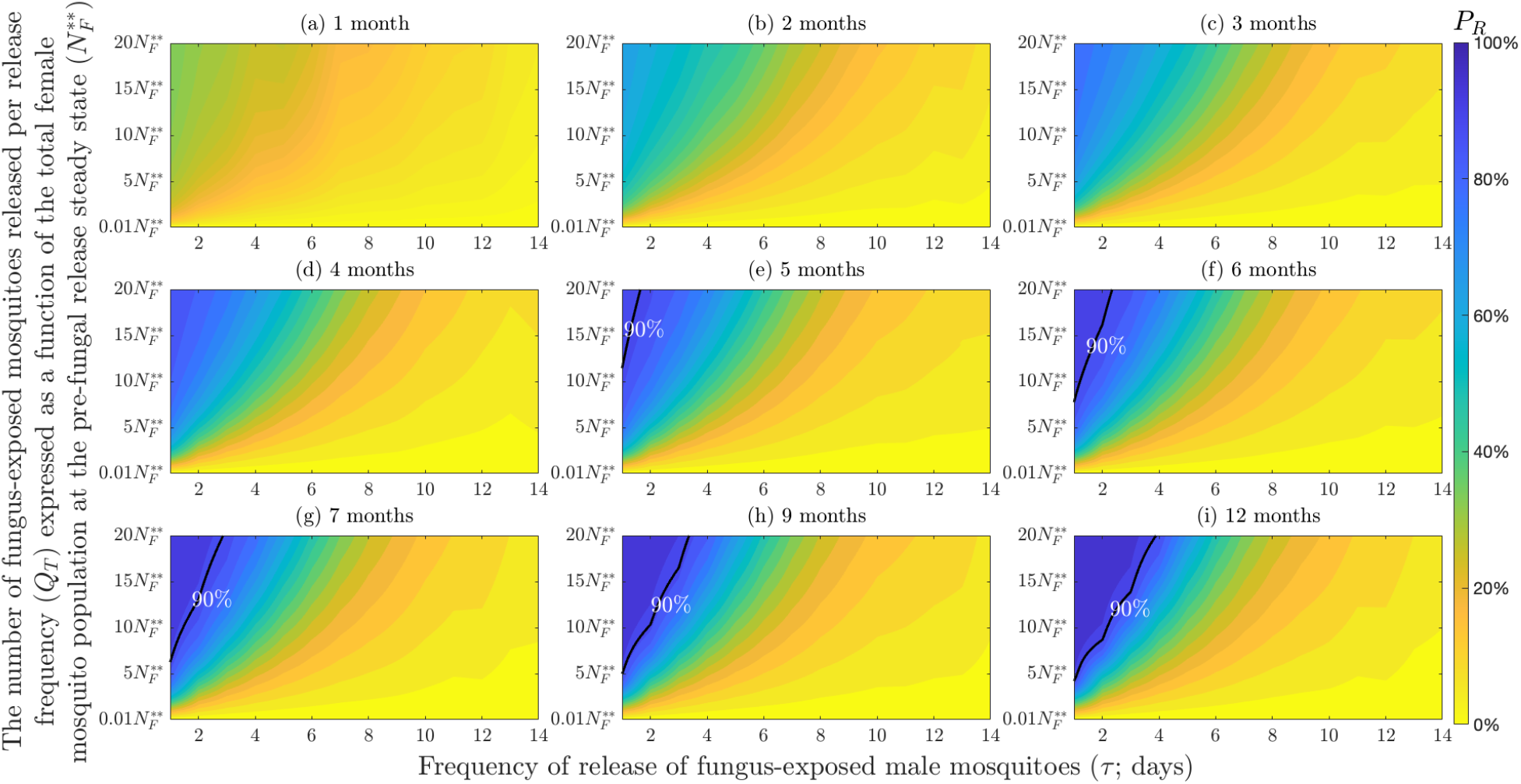
Contour plots of the percent reduction in the total population of adult female mosquitoes (*P*_*R*_, defined by (3.1)), as a function of the total number of male mosquitoes released *per* release period (*Q*_*T*_) and various values of release frequency (*τ*), corresponding to various durations of the male release program. The contour plots show *P*_*R*_ at (a) 1 month (b) 2 months (c) 3 months (d) 4 months (e) 5 months (f) 6 months (g) 7 months (h) 9 months and (i) 12 months from the initiation of the periodic fungus-exposed male mosquito release program. For simplicity, the release amount is given as a function of the total female population at steady state before fungus release 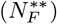. The black curve represents the level set where *P*_*R*_ = 90%. The baseline values of the parameters in Table 3 are used to generate these contour plots.

In addition to the logistical challenges outlined in Figure 7 regarding the fungus-exposed adult male release program, the program also faces issues related to the relatively short *rebound time*. The rebound time is formally defined as the minimum duration required for the female mosquito population to revert to its pre-fungal release steady-state 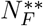, after the cessation of fungus-exposed adult male mosquito release program. Specifically, illustrated in Figure 8, this result depicts contour plots of the rebound time for the total adult female population to revert to their pre-male release steady-states, as a function of the total number of male mosquitoes released per release period (*Q*_*T*_) and different values of release frequency (*τ*), corresponding to various durations of the fungus-exposed male release program. This simulation demonstrates that extending the duration of the fungal release program can lead to a higher rebound time. However, this simulation clearly shows that even if the fungus-exposed male mosquito release program is maintained for a full year, the mosquito population will return to its pre-fungal release steady state approximately 2 months after the program ends.

**Figure 8:**
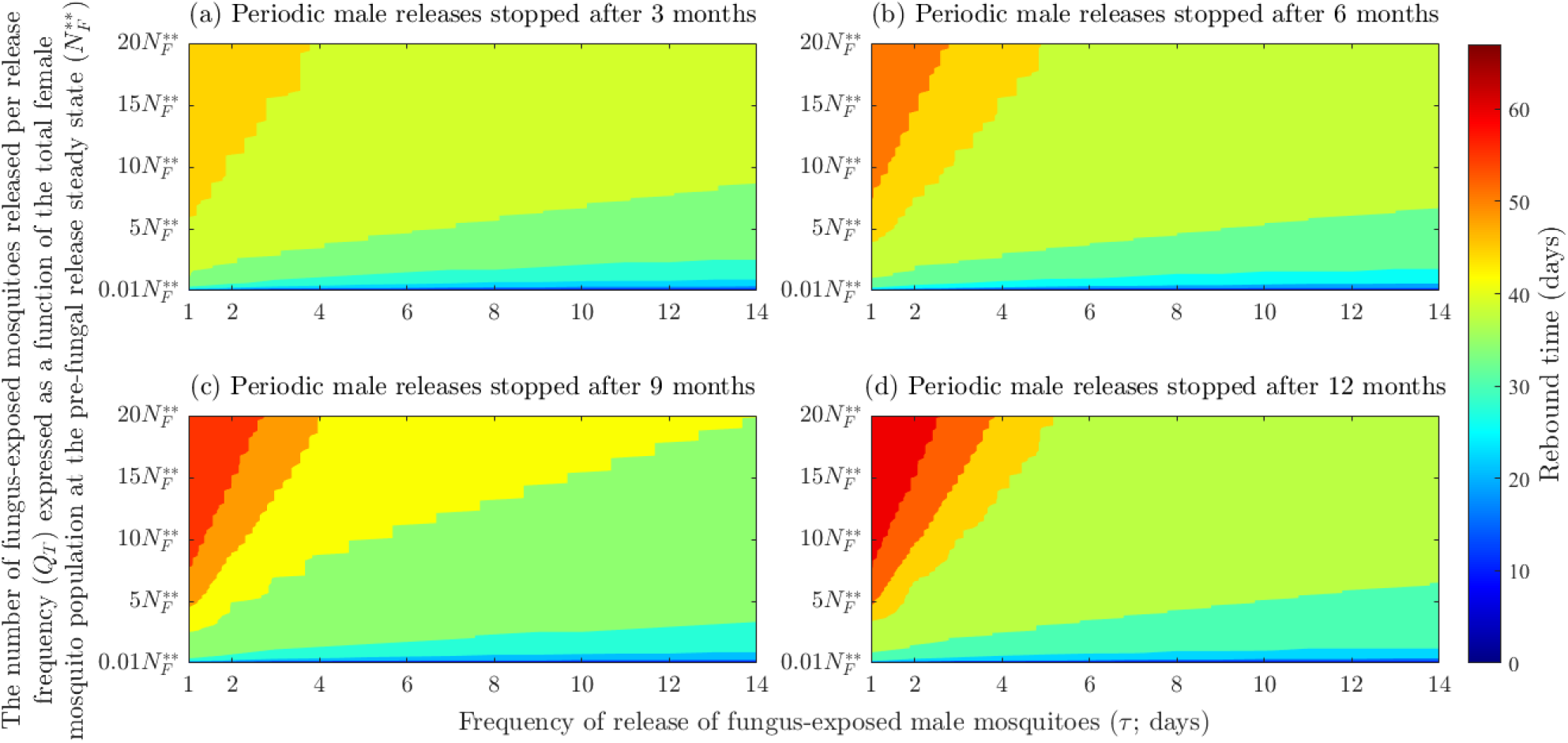
Contour plots of the rebound time for the total adult female population to revert to their pre-male release steady-states, as a function of the total number of male mosquitoes released per release period (*Q*_*T*_) and various values of release frequency (*τ*), corresponding to various durations of the fungus-exposed male release program. The rebound time is depicted for scenarios when fungus-exposed male mosquito release is stopped after (a) 3 months (b) 6 months (c) 9 months and (d) 12 months. The rebound time is calculated from the last day of release of fungus-exposed male mosquitoes. For simplicity, the release amount is given as a function of the total female population at steady state before the fungus release 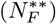. The remaining parameters are provided in Table 3.

## 4 Discussion and Conclusions

Owing to the widespread resistance of *Anopheles* mosquitoes to all the currently-available insecticides used in malaria endemic areas, several biocontrol methods have been proposed as alternatives to insecticide. These include sterile insect technology, the use of *Wolbachia*-based intervention, and the release of gene drives with desirable vector control characteristics [11–15]. Another recently proposed biocontrol method involves using a specific fungal species (*Metarhizium pingshaense*) to infect the male *Anopheles* mosquitoes [25, 26]. Mating between a male mosquito infected with the fungus and a wild female will result in transmission of the fungus to the female mosquito [25, 26]. The infected female will gradually sicken and eventually die from the fungal infection, thereby reducing the number of eggs laid and consequently, the number of new adult mosquitoes in the environment. This approach is one of several being trialed at a study site in Soumousso, a savanna town in Burkina Faso, targeting *Anopheles coluzzii*, one of the predominant malaria vectors in the area. Therefore, it is instructive to provide a mathematical modeling framework for assessing the population-level impact of this biocontrol method against malaria mosquitoes in the study area. To the best of our knowledge, our study is the first attempt to model and theoretically analyze the potential impact of a fungus (*M. pingshaense*) infection-based strategy to control the population abundance of malaria mosquitoes in a malaria-endemic setting. Specifically, our modeling study involves the design, analysis, and simulations of a novel mathematical model, which takes the form of a deterministic system of nonlinear differential equations. This model accounts for the transmission of the *M. pingshaense* fungus to uninfected adult female *Anopheles* mosquitoes through contact with fungus-infected male mosquitoes during mating and contact with mosquito cadavers carrying sporulating fungus. The model is analyzed for the case of both single and periodic releases of the fungus-infected male mosquitoes into the environment.

Rigorous analysis of the single release (baseline) model reveals that its non-trivial mosquito-present fungus-free equilibrium is locally-asymptomatically stable whenever the associated basic reproduction number (denoted by ℛ_0_) is less than one, and unstable when ℛ_0_ > 1. Ecologically, this means that a small influx of fungus-infected male mosquitoes will not generate a large number of fungus-infected mosquitoes males and females) in the environment when ℛ_0_ *<* 1. However, if ℛ_0_ > 1, even a small release of fungus-exposed male mosquitoes may allow the *M. pingshaense* fungus to establish itself within the mosquito population.

Since entomologists have not observed mosquito cadavers carrying sporulating fungus in semi-field experiments [25], we analyzed the baseline model for the case where fungal transmission to mosquitoes via contact with fungus sporulating cadavers does not occur (i.e., we set the parameter *β*_*c*_ = 0). In this case, our analysis revealed that the associated reproduction number of this special case of the baseline model 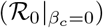 is precisely zero. This is because, without cadaver-mediated fungal transmission, fungus-infected male mosquitoes cannot generate new fungus-infected male mosquitoes through mating with fungus-uninfected females. Thus, this study shows that, in the absence of transmission from contact with fungus colonized cadavers (i.e., *β*_*c*_ = 0), the *M. pingshaense* fungus cannot independently survive or persist in the mosquito population over time. Overall, this theoretical finding suggests the necessity for external intervention, such as periodic releases of fungus-infected male mosquitoes, to enhance the potential effectiveness of the fungal infection-based mosquito control program. Furthermore, the associated reproduction number being zero emphasizes the importance of the use of other metrics, such as percentage reduction in female mosquito population during the fungus release program compared to pre-fungal release levels, to realistically quantify the population-level impact of the fungal-based mosquito control program. This is because, unlike in other ecological and/or epidemiological settings, where the reproduction number is often the main metric for measuring the persistence or lack thereof of a species or disease in a population, the associated basic reproduction number cannot be used to analyze the impact of the model parameters on these processes as it is zero.

The baseline model was extended to account for the periodic releases of adult male *Anopheles coluzzii* mosquitoes exposed to the transgenic *M. pingshaense* (specifically, Met-Hybrid strain) fungus. The simulations of the resulting periodic release model, which is in the form of an impulsive system of nonlinear differential equations, showed that a 90% reduction in the adult wild female mosquito population can only be achieved when the fungus-exposed male mosquitoes are released in high quantities (equivalent to around 12 times the steady-state value of the adult wild female population in the community) with high frequency (e.g., daily releases) for at least a five-month duration. However, this dramatic suppression of the wild mosquito population is short-lived, as the wild mosquito population rebounds to its pre-fungal intervention level within about two months after cessation of the fungus intervention program. This rebound (within about two months of cessation) also occurs even when the fungus-based program is implemented for a year, with daily releases of about 20 times the total size of the pre-release wild adult female population. Somewhat similar rapid rebounds can occur when chemical insecticide use is stopped, but insecticides are comparatively easy to apply, and mosquito populations have become much harder to control because of insecticide resistance. In contrast, frequent releases of large numbers of fungus-exposed male mosquitoes, for an extended period, may not be realistically attainable and sustainable in most malaria-endemic settings.

We compared the efficacy of the fungus-based biocontrol method to some other biocontrol methods for mosquito control. A modeling study by Smidler et al. [68] demonstrated that releasing genetically-modified adult male mosquitoes at a 10:1 release ratio (approximately 10 genetically-modified adult male mosquitoes for every wild adult mosquito in the target area) can achieve at least 90% suppression of the wild mosquito population over approximately 22 weeks of weekly releases. In contrast, the periodic fungal release program fails to achieve equivalent suppression levels within the same time frame, even with comparable release ratios of fungus-exposed mosquitoes. Conversely, Carvalho et al. [69] reported an 81% reduction of wild *Aedes aegypti* mosquito population (based on ovitrap indices) in Juazeiro, Brazil, by releasing a very large number of transgenic male mosquitoes (68.5 times the estimated number of the wild mosquito population in the community) three times every week for 6 months. In comparison, our fungus-based control can achieve 86% reduction in the wild mosquito population with similar release frequency (every three days) and much smaller release sizes (e.g., approximately 10:1 release ratio, which is equivalent to releasing 10 fungus-exposed male mosquitoes for every one wild mosquito in the pre-fungal program steady-state) for approximately six months. This suggests that the fungus-based method may outperform some other DNA-based biological methods and as such presents a viable alternative, particularly in settings where regulatory, ethical, or public acceptance concerns restrict the deployment of genetically-modified mosquitoes in the target area [70, 71].

Although this study provides crucial insights into the potential effectiveness and utility of fungus-based control of the malaria vector in a malaria-endemic setting, it has several limitations. For example, the study does not account for the seasonality of the mosquito population in malaria-endemic regions. Incorporating seasonal variations would enable the theoretical identification of the optimal deployment timing for the fungus-based interventions to maximize mosquito population reduction during malaria transmission seasons. Furthermore, because we assume that the mosquito population is constant throughout the year, we may be overestimating the release amount needed to achieve the 90% reduction in the adult wild female mosquito population and underestimating the time it takes for the female mosquito population to return to pre-fungal release steady state. Another significant limitation is the lack of explicit consideration of indoor fungus deployment methods in the modeling process. Indoor fungus application, a major strength of the fungus-based intervention, currently involves hanging black cotton sheets treated with fungal spores on household walls. When (male or female) mosquitoes encounter these sheets, they become infected, leading to their eventual fungus-induced death [24–26]. Other innovative application methods are being developed, including the use of baited traits that lure mosquitoes to the fungus. We aim to address these limitations and expand the scope of our model in future studies as it’s possible that simultaneous or alternating application of multiple mosquito control technologies could have additive or even synergistic effects. It is unlikely that a single method of control can provide a stand-alone intervention to tackle the rising emergence of mosquito vectors. However, using males as a vehicle to disseminate the fungus to mate seeking females would target malaria vector species that are not affected by IRS, bednets and fungus-impregnated sheets as well as outdoor biting mosquito populations that currently escape conventional control tools.

Given the well-documented resistance of *Anopheles* mosquitoes to long-lasting insecticidal nets (LLINs) [13], and the absence of observed resistance to pathogenic fungi in insects [72], fungus-based approaches in general are a promising alternative. Preliminary laboratory studies in Burkina Faso further demonstrate that *Anopheles* mosquitoes do not develop resistance to Met-Hybrid, making it logical to combine these two indoor approaches to tackle the challenges posed by LLIN resistance. We aim to extend this fungus-based modeling study to allow for the theoretical assessment of the combined fungus-based and LLINs-based control methods against mosquitoes in malaria endemic regions. Overall, this study, which provides the first comprehensive compartmental modeling framework for assessing the population-level impact of a fungus-based mosquito control intervention in a malaria-endemic setting, indicates that the prospect for significant reduction or suppression of the wild mosquito population is promising using the fungus-based program. Future studies could build upon the modeling framework and parameterization provided in this study to gain further insights to optimize the large-scale and widespread implementation of the fungus-based program to achieve maximum and sustained reduction of the malaria mosquito population in endemic areas.

## Acknowledgments

The authors thank the Brin Mathematics Research Center at the University of Maryland for funding the “Mathematics of Malaria Transmission Dynamics” workshop, where the impetus for this collaborative work was initiated (the visit of co-author EB was particularly instrumental to this effort). Etienne Bilgo is supported by the Wellcome Trust / National Institute for Health Research (NIHR), grant reference 218771/Z. ABG acknowledges the support, in part, of the National Science Foundation (Grant Number: DMS-2052363; transferred to DMS-2330801).

## A Model for the periodic releases of fungi-exposed male mosquitoes

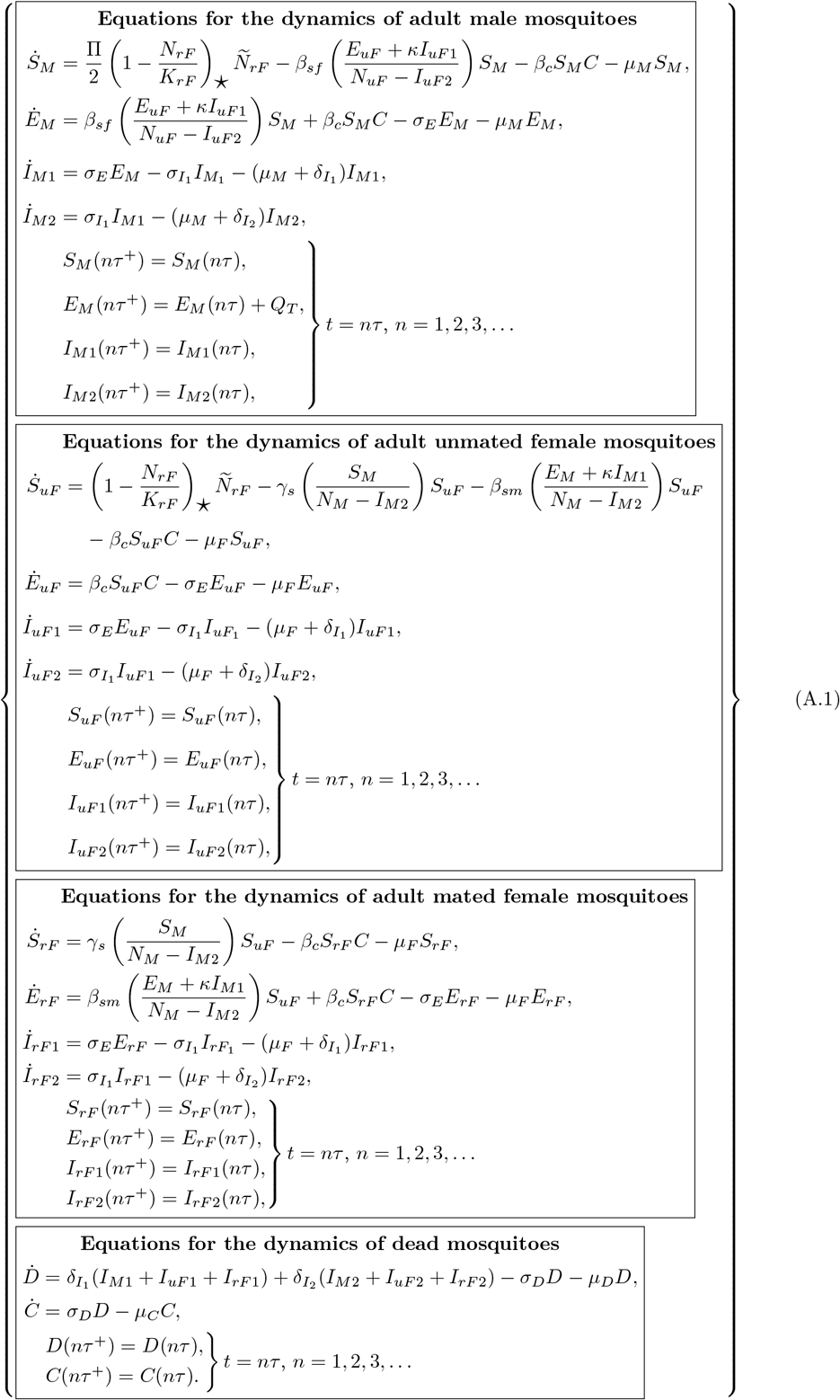

